# Comparative proteomic analysis of skin wound healing responses to biomaterial treatments identifies key pathways which govern differential regenerative outcomes

**DOI:** 10.1101/2025.05.19.654672

**Authors:** Alejandra Suarez-Arnedo, Eleanor L.P. Caston, Yining Liu, Hongxia Baia, David Muddiman, Tatiana Segura

## Abstract

The wound healing cascade is characterized by the steady progression of distinct stages. Though biomaterials are used clinically to enhance wound closure rate and quality of healed tissue, their mechanisms of action are less understood. Here we use proteomic analysis to characterize changes in the wound healing response across three biomaterial treatments: a clinically used collagen hydrogel, and two synthetic biomaterials that are characterized by an increased regenerative response either through decreased fibrosis or through an activation of adaptive immunity. We identified close to 5,000 proteins shared across the biomaterial treatment groups, sampled at timepoints representing the inflammation, proliferation, and resolution phases of wound healing. The collagen hydrogel maintains an enrichment of immune-related pathways throughout the healing process. The fibrosis-suppressing material enriches gene ontology (GO) terms related to increased epidermis development pathways, collagen synthesis, and collagen fibril organization. In contrast, the adaptive immunity-activating biomaterial shows an early enrichment of GO terms related to broad immunity and inflammation. Later, this same material promotes keratinization, muscle and lipid oxidation GO pathways. Taken together, this work determines the key temporal pathways (immunity, keratinization, muscle system process, and ECM organization) mediated by three biomaterials, which result in varying healed tissue structure.

## 1. Introduction

Dermal wound healing is a complex process that relies on the coordination of numerous biological pathways to restore the skin’s structure and function. Acute skin healing involves several overlapping phases, beginning with an early inflammatory response followed by cellular migration, proliferation, matrix deposition, tissue formation, and remodeling. This process is carefully programmed with an end goal of rapidly restoring the barrier function of skin, which in turn helps mitigate the risk of infection via harmful microbes and foreign material^[1]^. The body’s attempt to swiftly mend the wound favors rapid deposition of fibrotic granulation tissue, commonly known as scar^[2]^. Unfortunately, this resulting scar tissue lacks the structural integrity and functional dermal features of uninjured skin.

Materials for wound care date back thousands of years, with early civilizations using a variety of natural substances to treat and protect wounds, such as honey, plant-derived extracts, oils, and bandages soaked in wine or beer^[3]^. Even then, different materials were used for different types of wounds and achieved different levels of success. Modern materials for wound closure include biomaterials, which can simultaneously serve as delivery vehicles for therapeutics such as growth factors^[4]^, siRNA^[5]^, plasmid DNA^[6]^, and cells^[4a, 7]^, while also serving as a scaffold to support the growing tissue. However, these materials can also be engineered to promote wound closure and improve tissue deposition in the absence of therapeutic delivery, through engineering the cell-biomaterial interaction (e.g., mechanical properties, degradation rate, integrin binding, porosity, hydrophobicity). Examples of these materials include natural scaffolds such as fibrin^[8]^ and collagen^[9]^, and synthetic scaffolds including peptide hydrogels^[10]^, precast-porous hydrogels^[11]^, and microporous annealed particle (MAP) scaffolds^[12]^. While there have been promising results in the field of biomaterials for wound regeneration, studies aiming to understand the way in which materials with known beneficial outcomes reprogram the traditional healing pathways are lacking compared to the number of studies on biomaterial outcomes.

Proteomic technologies have had vast improvements in recent years, creating opportunities for in-depth studies on the effect of biomaterials in skin wound regeneration and beyond^[13]^. Proteomic analyses have been widely used in the wound healing field as an unbiased quantitative approach to provide insight into the complex biological mechanisms associated with the healing cascade^[14]^. Wound healing in both *Acomys cahirinus*, a mouse with unique skin regenerative capabilities, and *Mus musculus*, a mouse that heals via scarring, was investigated through proteomics on multiple time points^[15]^. Analysis of the differentially enriched proteins using gene ontology (GO) highlighted the proteasome and Wnt pathways, among others, for *Acomys*, whereas *Mus* enriched pathways included complement and extracellular matrix (ECM) interaction. Beyond this study, there have been additional proteomic investigations that aimed to uncover information that could be leveraged to improve wound healing, ranging from investigating the profiles of healing and non-healing wounds^[16]^ or cells isolated from them^[17]^, to creating a spatially resolved map of the skin proteome^[18]^. Other tissues have been investigated to better understand the influence of differential biomaterial treatments on healing, including regenerative responses to a titanium biomaterial bone implant^[19]^. Yet, to date, there has not been a proteomics study that investigates skin wound healing outcomes influenced by a biomaterial.

Thus, in this study, we aimed to investigate three biomaterial treatments with known differential outcomes in a dermal wound healing setting, from biomaterials that promote wound closure to those that promote regenerative healing. We designed a time-course study spanning the breadth of the wound healing response-including the different stages of healing-and compared tissue responses using proteomic analyses. Employing proteomic analyses of our conditions provided a comprehensive view of key proteins and pathways associated with biomaterial-mediated scarring or regenerative healing. Given the complexity of wound healing, we focused our investigations using GO analyses of biological pathways. Our study revealed a comprehensive understanding of the temporal interactions between biomaterial treatments and the skin. The data presented here provide valuable insight into pathways that can be temporally targeted to modulate wound resolution outcomes.

## 2. Results and Discussion

### 2.1 The wound healing cascade is visible through its proteome

Transcriptomics is widely used to study protein expression changes as biological processes are taking place^[20]^; however, gene expression is not equivalent to protein expression. Numerous studies have shown that mRNA transcripts are not always translated to protein, and largely depend on context such as alternative splicing and post-transcriptional regulation^[21]^. In contrast, proteomics measures protein expression directly, allowing for a clear understanding of protein levels and their biological context. Thus, we chose to use proteomics to understand the diverse mechanisms by which biomaterials promote the healing of skin wounds. We studied three biomaterials with distinct healing outcomes: Woun’Dres®, a clinically used collagen based hydrogel that promotes a moist wound environment and faster clearance of dead tissue^[22]^; LMAP, a synthetic hydrogel that was previously found to accelerate wound closure, lower inflammation, and reduce fibrosis^[12a, 23]^; DMAP, a synthetic hydrogel with a D-amino acid (D-AA) peptide crosslinker that was previously found to induce regenerative healing through an adaptive immune response^[24]^. LMAP and DMAP were synthesized as previously described^[24]^ and were matched in mechanical properties and hydrogel microparticle size (**Supplemental Figure 1A-E**), while Woun’Dres® was used as supplied by the manufacturer.

Full thickness wounds were created in the backs of hairless mice using a circular biopsy punch (5-mm), and the wound bed was treated with each biomaterial (10µL) as a flowable liquid. MAP biomaterials undergo a gel transition and become solids at 30 minutes post-injection, while Woun’Dres® maintains its viscous liquid state **(Supplemental Figure 1F)**. The wound morphological outcomes of the chosen biomaterials have been previously established^[25]^. Treatment with Woun’Dres® resulted in a scar-like phenotype with less collagen regeneration (reduced collagen area) and increased fibrosis compared to MAP scaffolds^[23]^. Treatment with LMAP promotes a less scar-like phenotype compared to Woun’Dres®, with improved ECM quality (reduced αSMA+ area, increased collagen area) and less inflammation (reduced CD11b+ area) in the wound bed^[23]^. DMAP treatment encouraged *de novo* regeneration as characterized by physiological undulation of the epidermis and increased hair follicles per section, which features a prominent SOX9+ bugle stem cell region^[24]^. Histological characterization of wound sections at the study endpoint (Day 21) confirmed the previously observed healing outcomes for each biomaterial (**Figure 1A**, **Supplemental Figure 2**). Treatment with DMAP resulted in increased regeneration of hair follicles. Quantification showed a higher density of hair follicles per millimeter of wound tissue and a reduced afollicular percentage (indicative of true scar region) in DMAP-treated wounds compared to the Woun’Dres® control, which was not observed in LMAP wounds **(Supplemental Figure 2A-E)**.

**Figure 1.**
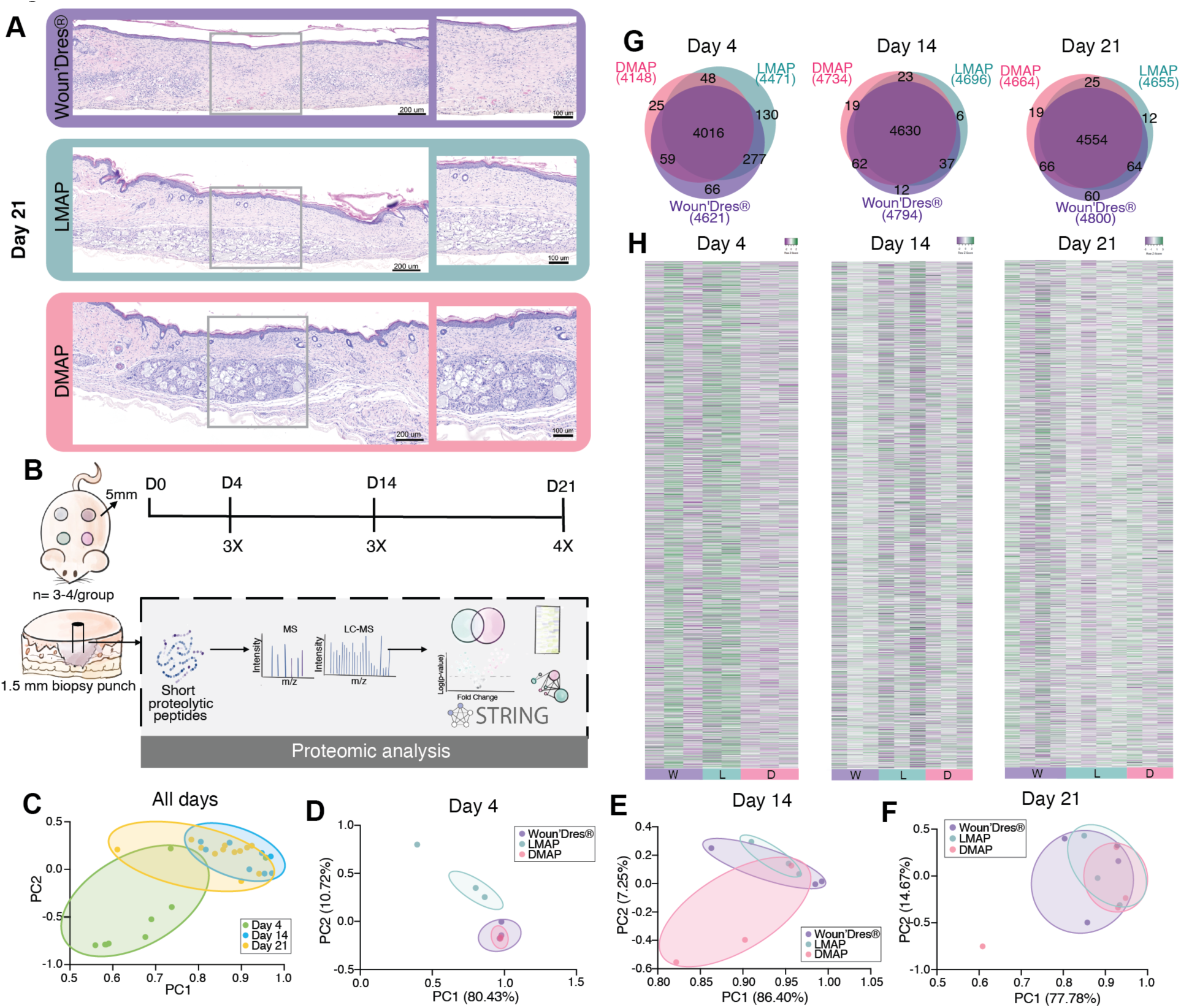
Experimental design and proteomic analysis of wounds treated with DMAP, LMAP or Woun’Dres®. **A)** Histological analysis of wound healing at day 21: Representative histological sections stained with hematoxylin and eosin (H&E) show differential wound healing capabilities for three different treatment groups. Scale bars represent 200 μm. **B)** Schematic of the experimental timeline and sample Collection: A 5 mm wound was created on the dorsal skin of mice (n=3-4/group/day), followed by collection of 1.5 mm biopsy punches at days 0, 4, 14, and 21 post-wounding. Proteomic analysis was conducted using mass spectrometry (MS) and liquid chromatography-mass spectrometry (LC-MS), with data analyzed for protein interactions using STRING V11. **C)** Principal Component Analysis (PCA) plots illustrate the grouping of samples based on proteomic profiles at different time points. **D-F)** PCA plots groupings of samples, pooled by biomaterial treatment at day 4, 14 and 21. **G)** Venn diagrams of protein overlap display the number of unique and shared proteins identified in each treatment group at days 4, 14, and 21. **H)** Heatmaps show the expression levels of proteins across different treatment groups at each time point. Rows represent individual proteins, while columns represent replica samples from each treatment group. Color gradients indicate relative expression levels.

To capture the proteome throughout the wound healing cascade, we harvested tissue using a 1.5 mm biopsy punch at the wound center on Day 4 (inflammation phase), Day 14 (proliferative phase), and Day 21 (remodeling phase) for bottom-up proteomics **(Figure 1B).** Liquid chromatography-tandem mass spectrometry (LC-MS/MS) was performed following sample homogenization and a modified filter-aided sample preparation (FASP) ^[26]^. Raw data was mapped against the *Mus Musculus* FastA database.

We first performed principal component analysis (PCA) to find overarching differences between the samples. As expected, the proteome of each stage of healing is distinct from each other, regardless of biomaterial treatments **(Figure 1C)**, indicated that the healing cascade is largely the same for each biomaterial. PCA analysis revealed that Day 4 samples are distinct from Days 14 and 21, as indicated by their non-overlapping distribution along the principal components. In contrast, Day 14 and 21 samples show overlapping distributions in the PCA space, corroborating that the proteome of the early wound healing response (Day 4) is unique compared to later stages. PCA executed by day *and* biomaterial treatment highlighted Day 4 as the most compact distribution within each biomaterial treatment group **(Figure 1D)**, illustrating that the proteome of the wound environment is spatially similar early during wound healing. Although the wound tissue for proteomic analysis was obtained from similar regions, we observed increased variability within treatment groups as wound healing progressed **(Figure 1E, F)**, suggesting the wound healing response is not uniform throughout the wound. DMAP and Woun’Dres® PCA points are close together early on (Day 4) but diverged at later time points (Day 14, 21) **(Figure 1D-F)**. The Pearson R correlations align with the PCA observations, with correlations ranging from 0.95-1 on Day 4, to 0.54-0.95 and 0.27-0.96 on Days 14 and 21, respectively (**Supplemental Figure 3A-F**).

The proteome LC-MS/MS data was then log transformed and normalized to create a normal distribution data set with reduced bias and improved comparability^[27]^. An outlier detection method was applied to identify highly variant biological samples using hierarchical clustering with average linkage and Euclidean distance. This resulted in the removal of one LMAP replicate on Day 4 and one DMAP replicate on Day 21 **(Supplemental Figure 3G-I)**.

Proteomic data processing yielded the identification of nearly 5,000 unique proteins. Of these proteins, there was substantial overlap, with 4,106 (87%), 4,630 (97%), and 4,554 (95%) detected in all groups on Days 4, 14, and 21, respectively **(Figure 1G, H)**. Across all timepoints, Woun’Dres®, LMAP, and DMAP expressed a cumulative of 133, 146, and 61 unique proteins. Interestingly, 2 proteins (Fem1ab, Dtx4) were found to be unique to wounds treated with MAP biomaterials, with exclusive expression in DMAP on Day 14 and exclusive expression in LMAP on Day 21 **(Supplemental Data 1-3)**. These proteins are linked to ubiquitin ligase activity which has been shown to be important in regulating the removal of damaged proteins as well as cell migration and proliferation in wound healing^[28]^. Heatmaps of the proteome illustrate the expanse of proteins identified and their range of expression across treatment, replicate, and timepoint **(Figure 1H)**. The heatmaps corroborate the trends identified with the PCA; Day 4 exhibits the most consistent expression patterns across treatment groups and replicates.

Because proteomic analysis best compares two conditions at a time, our subsequent analysis is presented as two-way comparisons at each time point –Woun’Dres® versus LMAP (Comparison 1), LMAP versus DMAP (Comparison 2), Woun’Dres® versus DMAP (Comparison 3)– to identify upregulated proteins using volcano plots and Gene ontology (GO) enrichment analysis. Heatmaps of proteins identified via non-GO analyses are represented in purple and green, whereas heatmaps of proteins identified via GO analyses are represented in blue and red. All plots incorporate all three biomaterial treatment groups, but an asterisk and comparison symbol are used to highlight which group was not included in the two-way analysis.

### 2.2 LMAP wound treatment results in the most differentially expressed proteins on Day 4

Volcano plots of differentially expressed proteins on Day 4 for each comparison highlighted the early differences in skin healing response to the biomaterial treatments **(Figure 2A-C)**. Of the 4,646 proteins found on Day 4, Woun’Dres® versus LMAP had 74 and 267 upregulated proteins, LMAP versus DMAP had 1,024 and 235 upregulated proteins, and Woun’Dres® versus DMAP had 326 and 172 upregulated proteins. Thus, different biomaterial treatments can elicit different protein expression profiles on the same animal.

**Figure 2.**
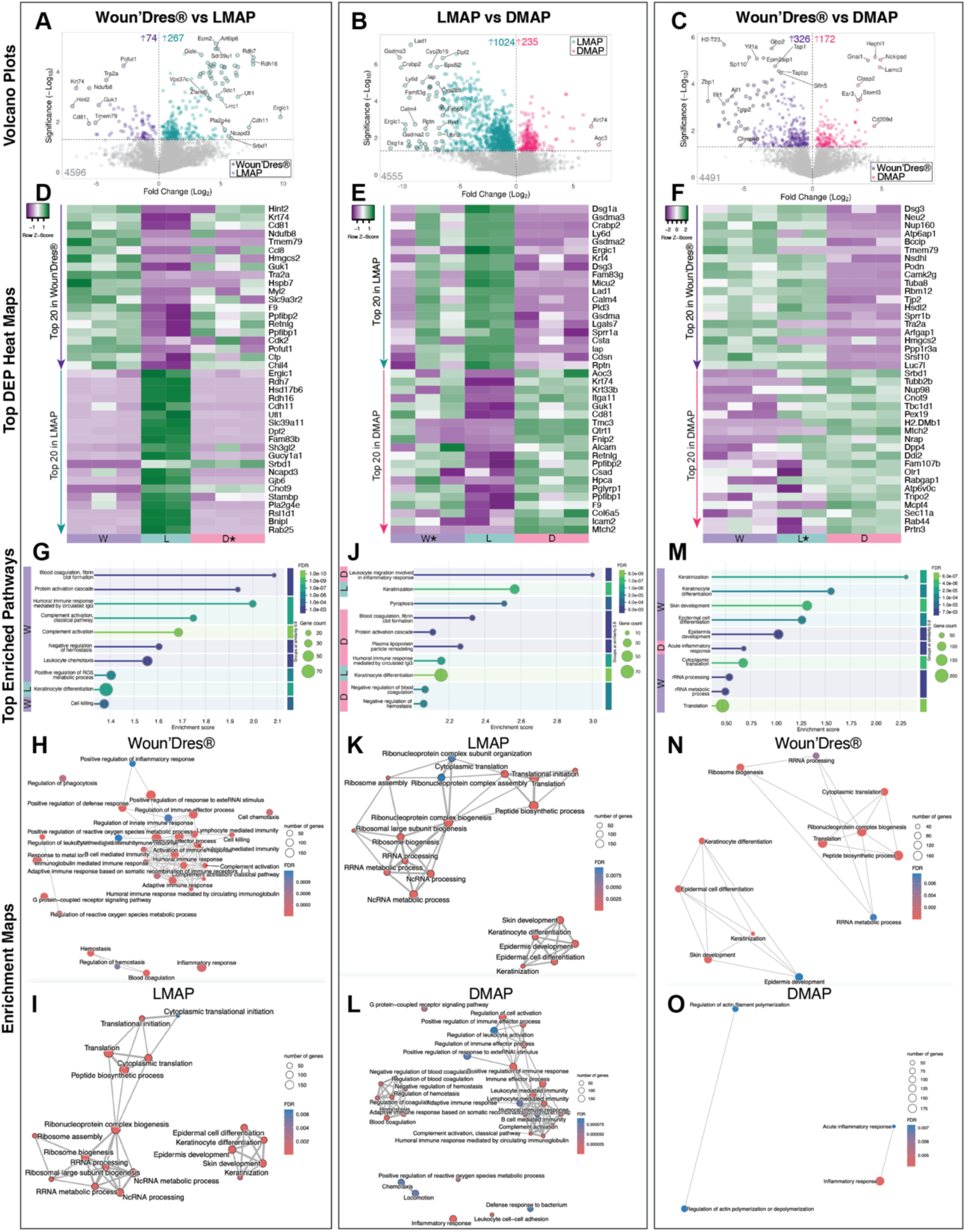
Proteomic analysis at day 4. **A-C)** Volcano plot showing the fold change (Log_2_) versus significance (-Log_10_ p-value) for proteins in **(A)** Woun’Dres® vs LMAP, **(B)** LMAP vs DMAP, and **(C)** Woun’Dres® vs DMAP. **D-F)** Heatmaps show the expression levels of the 20 most upregulated proteins for each treatment in descending order for all three comparisons. **G)** Gene ontology enrichment analysis comparing biological processes between Woun’Dres® and LMAP, highlighting 10 of the most significantly enriched pathways **H-I)** Enrichment GO pathways for comparison 1 of (**H)** Woun’Dres® and (**I)** LMAP are indicated by node size (representing the number of genes) and color (indicating false discovery rate - FDR). **J)** Gene ontology enrichment analysis comparing biological processes between LMAP and DMAP, highlighting 10 of the most significantly enriched pathways. **K-L)** Enrichment GO pathways comparing LMAP and DMAP of **(K)** LMAP and **(L)** DMAP are indicated by node size (representing the number of genes) and color (indicating false discovery rate - FDR). **M)** Gene ontology enrichment analysis comparing biological processes between Woun’Dres® and DMAP, highlighting 10 of the most significantly enriched pathways. **N-O)** Enrichment GO pathways comparing Woun’Dres® and DMAP of **(N)** Woun’Dres® and **(O)** DMAP are indicated by node size (representing the number of genes) and color (indicating false discovery rate - FDR).

With the significant differentially expressed proteins (DEP, P<0.05) highlighted by the volcano plots for each comparison, we plotted the top 20 for each condition using heat maps in descending order to further visualize the expression patterns across biological replicates **(Figure 2D-F)**. As seen with the Day 4 PCA plots, in Comparisons 1 and 2, DMAP expression profiles aligned more closely with Woun’Dres® than LMAP **(Figure 2D-E).** The heatmaps assisted in identifying interesting protein expression similarities and differences between the three biomaterial treatments **(Supplemental Figure 4A-F).**

One family of proteins that is important in wound healing and whose comparative expression profile varied across treatments was the Keratin family. Woun’Dres® treatment upregulated Krt74, LMAP treatment upregulated Krt4, and DMAP treatment upregulated both Krt74 and Krt33b **(Figure 2D-E**, **Supplemental Figure 4A-C).** While keratins are important in wound healing^[29]^, Krt74 and Krt33b are more traditionally associated with the inner root sheath^[30]^ and hair structural components^[31]^, while Krt4 is known for its role in non-keratinized epithelial tissues^[32]^. Indicating that the biomaterials are differentially modulating non-traditional keratin pathways at this early timepoint.

Tmem79, a protein known to be essential in skin barrier function^[33]^, was strongly upregulated by Woun’Dres® in both Comparisons 1 and 3 **(Figure 2D, F**, **Supplemental Figure 4D).** Comparison 1 highlighted the upregulation of the Retinol dehydrogenase (Rdh) family by LMAP, with both Rdh7 and Rdh16 upregulated **(Figure 2D**, **Supplemental Figure 4E)**. Rdh16 is involved in retinol (a form of vitamin A) metabolism, with deficiencies in the vitamin A family known to negatively impair wound closure and collagen synthesis^[34]^. Within Comparison 2, the Desmogleins (Dsg) (Dsga1 and Dsg3) family was consistently upregulated by LMAP and downregulated by DMAP **(Figure 2E**, **Supplemental Figure 4F).** This family is comprised of cadherin-type transmembrane adhesion molecules essential for maintaining structural integrity^[35]^. Interestingly, these proteins are typically downregulated at the beginning of wound healing to allow for keratinocyte migration^[35]^. LMAP treatment resulted in the consistent upregulation of protein families tightly related to the ECM, whereas DMAP treatment downregulated these same proteins. Overall, these findings suggests that LMAP promotes an early wound healing response focused on enhancing structural integrity and increasing barrier function.

### 2.3 Day 4 comparative Gene Ontology analysis highlights biomaterial dependent mechanisms

Next, we employed GO analysis, focusing on functional enrichment visualization, enrichment maps, and hierarchal clustering to provide biological context to the common proteome identified. Functional enrichment visualization highlights the top 10 biological processes enriched with a false discovery rate (FDR) below 0.05 for each comparison. The enrichment score is an essential metric which quantifies how much the values associated with a specific term deviate from the dataset’s average^[36]^. Enrichment maps allow for a visual summary of GO pathway functional enrichment, along with the advantage of illustrating the relationships within the network^[36a]^. Hierarchal clustering provides complimentary analysis which focuses on network-based clustering beyond those that may be traditionally annotated^[36a]^.

A total of 74, 96, and 16 differentially expressed GO biological processes were identified for Comparisons 1, 2, and 3, respectively **(Supplemental Data 4-6)**. Comparison 1 highlighted the differences between Woun’Dres® and the baseline MAP treatment, LMAP **(Figure 2G-I)**. Woun’Dres® exhibited terms related to hemostasis, inflammation, the adaptive immune response and complement activation **(Figure 2G-H**, **Supplemental Figure 4G)**. In contrast, LMAP treatment enriched GO biological processes associated with skin development and epidermal cell differentiation in the early stages of wound healing **(Figure 2G, I**, **Supplemental Figure 4H)**. LMAP also had an enrichment of terms including ‘translation’, ‘ribosome assembly’, and ‘peptide biosynthetic process’ that were grouped by the enrichment tree as peptide and biosynthetic translation **(Figure 2D-E**, **Supplemental Figure 4H)**. Comparison 2 reinforced the LMAP pathways seen in Comparison 1, with repeated presence of pathways related to epidermal development and peptide synthesis **(Figure 2J-K**, **Supplemental Figure 4I).** This result when compared against DMAP was surprising, given the downregulation of the Dsg3 protein by DMAP, which is the expected expression pattern during early wound healing for re-epithelization and keratinocyte migration^[35]^.

DMAP had 74 enriched biological processes compared to LMAP’s 19 in Comparison 2 **(Supplemental Data 5)**. Of these, 40% were immune-related, all of which were upregulated in DMAP, including the top enriched term of ‘Leukocyte migration involved in inflammatory response’ **(Figure 2E,J,L)**. Previously we had found that DMAP treated wounds engaged the adaptive immune system to promote regenerative healing; mice lacking mature B or T cells treated with DMAP showed scar formation, while wild type mice showed regeneration^[25b]^. The enrichment trees also identified groupings related to response to bacterium **(Supplemental Figure 4J)**. Bacteria are known to produce a variety of D-AAs^[37]^, which play important roles in bacterial cell wall biogenesis, biofilm integrity, and spore germination^[38]^, providing interesting insight to how the body might view the peptide crosslinker containing D-AAs. From this, wounds treated with DMAP showed a proteome that had increased GO terms associated with a broad immune response and hemostasis compared to LMAP, closer to what was observed with Woun’Dres®.

Comparison 3 had the least number of GO biological processes identified (**Supplemental Data 6**), which aligned with our PCA observations for Day 4 (**Figure 1D**) that these two conditions are similar. When compared against DMAP, Woun’Dres® enriched ‘Keratinocyte differentiation’ and other epidermal development related terms, instead of the immune related terms seen in the Woun’Dres® profile in Comparison 1 **(Figure 2M-N**, **Supplemental Figure 4K).** DMAP maintained the decrease of epidermal development terms and enrichment of immune terms, as seen by the presence of ‘Acute inflammatory response’ within the top 10 enriched pathways **(Figure 2M, O)**. An enrichment tree was unable to be generated for DMAP **(Supplemental Figure 4L),** further illustrating non-significant differences in GO terms for proteins isolated from Woun’Dres® and DMAP treated wounds at this timepoint.

Day 4 is a critical juncture in the wound healing cascade, denoting the transition away from the inflammatory phase towards proliferative phase. These results suggest that LMAP facilitates an earlier transition from the inflammation phase towards the proliferative phase of healing as marked by a decrease in immune terms and an increase in peptide synthesis and keratinocyte differentiation. In contrast, both DMAP and Woun’Dres® appear to encourage an increase in immune response, although their specific roles differ. DMAP emphasizes actin filament regulation and inflammatory response, suggests a particular effectiveness in enhancing cell migration and managing inflammation during wound healing. On the other hand, Woun’Dres® focuses more on pathways related to protein synthesis and epidermal development, indicating its role in supporting the structural rebuilding of the skin.

### 2.4 LMAP treatment rapidly induces the expression of collagen and keratinocyte related proteins on Day 4

We applied the Matrisome bioinformatics approach to our proteomics datasets to gain deeper insights into the specific categories of ECM-related proteins, given the essential role the ECM plays in wound healing and tissue quality^[39]^. By Day 4, there were no appreciable differences in the core matrisome (collagens, ECM glycoproteins, proteoglycans) between the groups **(Figure 3A-B).** LMAP treatment appeared to have more matrisome-associated protein expression **(Figure 3A)**. When this category was further broken down into its components (ECM-affiliated proteins, ECM regulators, and secreted factors), a significant increase in LMAP was observed for ECM-affiliated proteins compared to DMAP **(Figure 3B).** Upon further analysis of the matrisome-associated groupings, ECM regulators was highlighted as the biggest contributor to the increase in matrisome-associated proteins for all groups **(Figure 3B)**. This finding aligns with the important role that ECM remodeling enzymes (transglutaminases^[40]^, MMPs^[41]^) and their regulators play in orchestrating the removal of damaged tissue and formation of new ECM components, processes especially prevalent during early stage wound healing.

**Figure 3.**
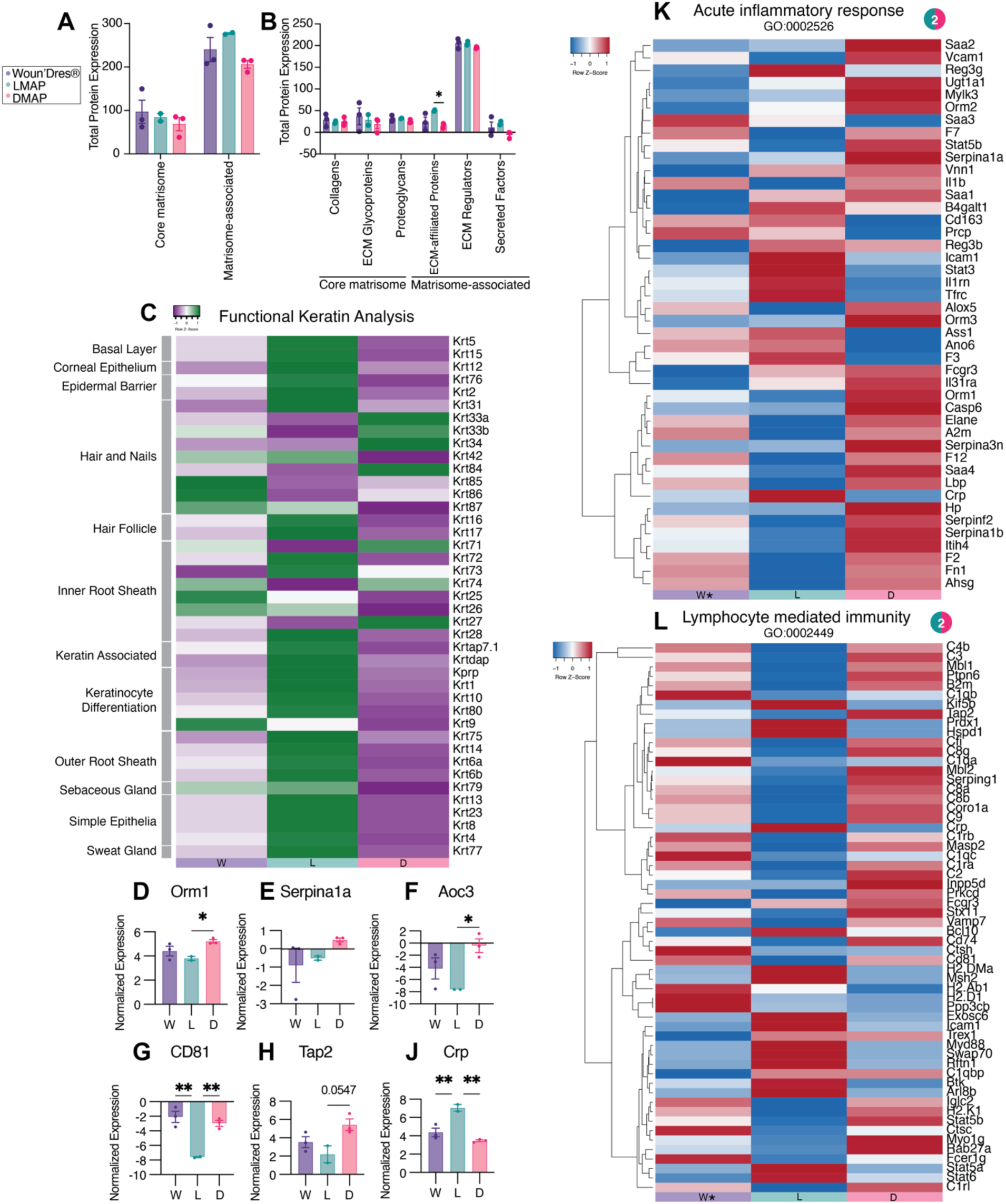
Detailed proteomic analysis of wound healing at day 4. **A)** Total protein expression in Core Matrisome and Matrisome-Associated Categories. Bar plots show total protein expression levels in the core matrisome and matrisome-associated proteins across three experimental groups: Woun’Dres® (W), LMAP (L), and DMAP (D). Two-way ANOVA was completed with Tukey’s multiple comparisons test (simple effects within rows). Significant differences are indicated (*p < 0.05). **B) P**rotein subcategories in Matrisome analysis. Detailed breakdown of protein expression within subcategories of the matrisome, including collagens, ECM glycoproteins, proteoglycans, ECM-affiliated proteins (proteins that share some architectural similarities with ECM proteins or that are known to be associated with ECM proteins), ECM regulators, and secreted factors. Two-way ANOVA was completed with Tukey’s multiple comparisons test (simple effects within rows). Significant differences between groups are marked (*p < 0.05). **C)** Heatmap illustrating keratin gene expression associated with various skin structures and functions, such as the basal layer, corneal epithelium, epidermal barrier, hair follicles, and sebaceous glands. Expression levels are normalized and color-coded (green: low expression; purple: high expression**). D-J).** Normalized expression of key proteins. Bar plots display normalized expression levels of specific proteins involved in inflammation and immunity**: (D)** Orm1 (acute phase protein) **(E)** Serpina1a (serine protease inhibitor), (**F)** Aoc (amine oxidase), **(G)** CD81 (immune cell marker), **(H)** Tap2 (antigen presentation), **(J)** Crp (C-reactive protein). One-way ANOVA with Tukey’s multiple comparisons test was performed. Statistical significance is indicated (*p < 0.05; **p < 0.01). **K)** Acute Inflammatory Response (GO:0002526) Heatmap showing differential protein expression related to acute inflammatory responses. Genes are hierarchically clustered, with red indicating upregulation and blue indicating downregulation across the experimental groups **L)** Lymphocyte-Mediated Immunity (GO:0002449) Heatmap depicting protein expression changes associated with lymphocyte-mediated immune responses. Red represents upregulated proteins, while blue indicates downregulated proteins.

Keratinocytes, the main epidermal cell type in the skin, are known to express a variety of keratins^[42]^. Given the hair follicle regenerative phenotype initiated by DMAP, we were curious to see how the proteins related to keratins varied over time. In GO biological pathway analysis, we observed that DMAP enriched immune related terms over those related to epidermal differentiation **(Figure 2J-L, 2M-O)**. Using data gathered from literature on the roles of different keratins^[18, 43]^, we organized keratins according to their functions based on treatment and day **(Figure 3C).** LMAP biomaterial treatment resulted in the greatest upregulation of keratin proteins across classifications ranging from keratinocyte differentiation to hair follicle **(Figure 3C)**. Interestingly, DMAP and Woun’Dres® had a stronger presence of hard (hair and nail) keratins, an unexpected result at this early time point **(Figure 3C).**

### 2.5 Day 4 immune activation is a critical mechanistic component of DMAP-driven tissue regeneration

To further elucidate the mechanistic underpinnings of the immune upregulation seen with Woun’Dres® and DMAP but not LMAP, we conducted comprehensive heatmap analyses of proteins found within specific GO terms, providing a high-resolution view of the immunological landscape. Compared to LMAP, both Woun’Dres® and DMAP promoted a broad and early immune response **(Figure 2)**. We chose to investigate heatmaps for a term from each component of the spectrum encompassed by the innate response and adaptive response **(Figure 3D-L)**.

A clear upregulation of proteins within the ‘Acute inflammatory response’ pathway was detected in DMAP **(Figure 3D-G, K)**. This included proteins within the Orm (Orm1, Orm2, Orm3) and Serpin (Serpina1a, Serpina3n, Serpinf2, Serpina1b) families. The Orm family of proteins are membrane proteins found within the endoplasmic reticulum that act as negative regulators of sphingolipid synthesis^[44]^. In the skin, sphingolipids (which include ceramides) play an important role in the epidermis barrier function and keratinocyte cell cycle^[45]^. In the immune system, regulation of sphingolipid homeostasis is important for immune cell polarization^[44]^. On Day 4, Orm1, which has been specifically shown to stimulate Interlukin-10 levels in monocytes for supporting an M2 pro-regenerative phenotype^[46]^ was significantly upregulated in DMAP when compared to LMAP **(Figure 3D)**. Despite seeing similar proteome profiles between DMAP and Woun’Dres® (**Figure 1D**), this significant upregulation was not seen in Woun’Dres®. Similarly, trends of high expression in DMAP were observed for Serpina1a **(Figure 3E)**, a protein within the Serpin family that regulates immune responses by acting as a serine protease inhibitor^[47]^. The Serpina1a protein is most attributed to inhibiting neutrophil elastase^[48]^. Serpina1 is one of the serpins found to be downregulated in diabetic mice (which suffer from chronically inflamed wounds), and delivery of Serpina1-loaded EVs resulted in faster wound closure in diabetic mouse wounds^[49]^. We also found that Aoc3, a protein found within the broader ‘Inflammatory response’ category, was significantly upregulated in DMAP, and was the top DEP against LMAP **(Figure 2E**, **Figure 3F**, **Supplemental Figure 5A).** Aoc3 is an essential protein expressed mainly on endothelial cell surfaces that helps to mediate the transmigration and recruitment of leukocytes into injured tissues^[50]^. The increase in proteins related to cell recruitment correlates well with histology data from Day 7 wounds that showed DMAP promoted more total cell recruitment to the infiltrating edge compared to an untreated wound **(Supplemental Figure 5B-E).**

Previously, the adaptive immune system was implicated in DMAP-mediated regeneration through both RAG1^-/-^ models and D-MMP-specific antibodies^[25b]^. Both DMAP and Woun’Dres® showed overlap in profiles for proteins within ‘Lymphocyte mediated immunity’, as seen by expression of proteins like CD81 **(Figure 3G, L).** This protein is required for normal B cell function and influences T cell activation^[51]^. Additionally, expression is linked to Th2 immune responses via IL-4 synthesis by lymphocytes^[52]^. Tap2 showed the highest expression in DMAP, indicating the upregulation of proteins that support CD8+ T cell function at Day 4 not seen in other groups **(Figure 3H, L).** Within this GO pathway, we observed overlap for many complement related proteins, which were also upregulated by Woun’Dres® and DMAP **(Figure 3L**, **Supplemental Figure 5F)**. These findings suggest that by Day 4 DMAP activates multiple immune-related pathways, often similar to those activated by Woun’Dres, but with a distinct profile characterized by a robust acute inflammatory response and enhanced immune cell recruitment.

Interestingly, C-reactive protein (Crp) stood out as one of the few proteins upregulated in LMAP treated wounds in both the ‘Lymphocyte mediated immunity’ and ‘Complement activation’ GO pathways **(Figure 3J, L**, **Supplemental Figure 5F)**. Crp, an acute phase protein, is known to increase during an injury response^[53]^ and has been shown to promote keratinocyte migration in *in vitro* scratch assays^[53]^. The downregulation of Crp by DMAP aligns with the observed early suppression of keratinocyte-related processes.

### 2.6 Proteome during the proliferative phase (Day 14) highlights ECM and immune proteins

As wound healing progresses, skin works to replace the provisional matrix first initiated by the fibrin clot with ECM deposited to form new granulation tissue^[54]^. Thus, the expectation for this timepoint is that the inflammatory stage would subside, and the proliferative/ECM deposition stage of wound healing would be active. Volcano plots were used to visualize significantly upregulated proteins within each group **(Figure 4A-C)**. The top 20 DEPs were once again highlighted using heatmaps for each comparison **(Figure 4D-F)**. While we surprisingly saw overlap of the Day 4 Woun’Dres® and DMAP proteomic profiles, this similarity diminished by Day 14, as noted by the increase in DEP expression intensity **(Figure 2F**, **4F)**.

**Figure 4.**
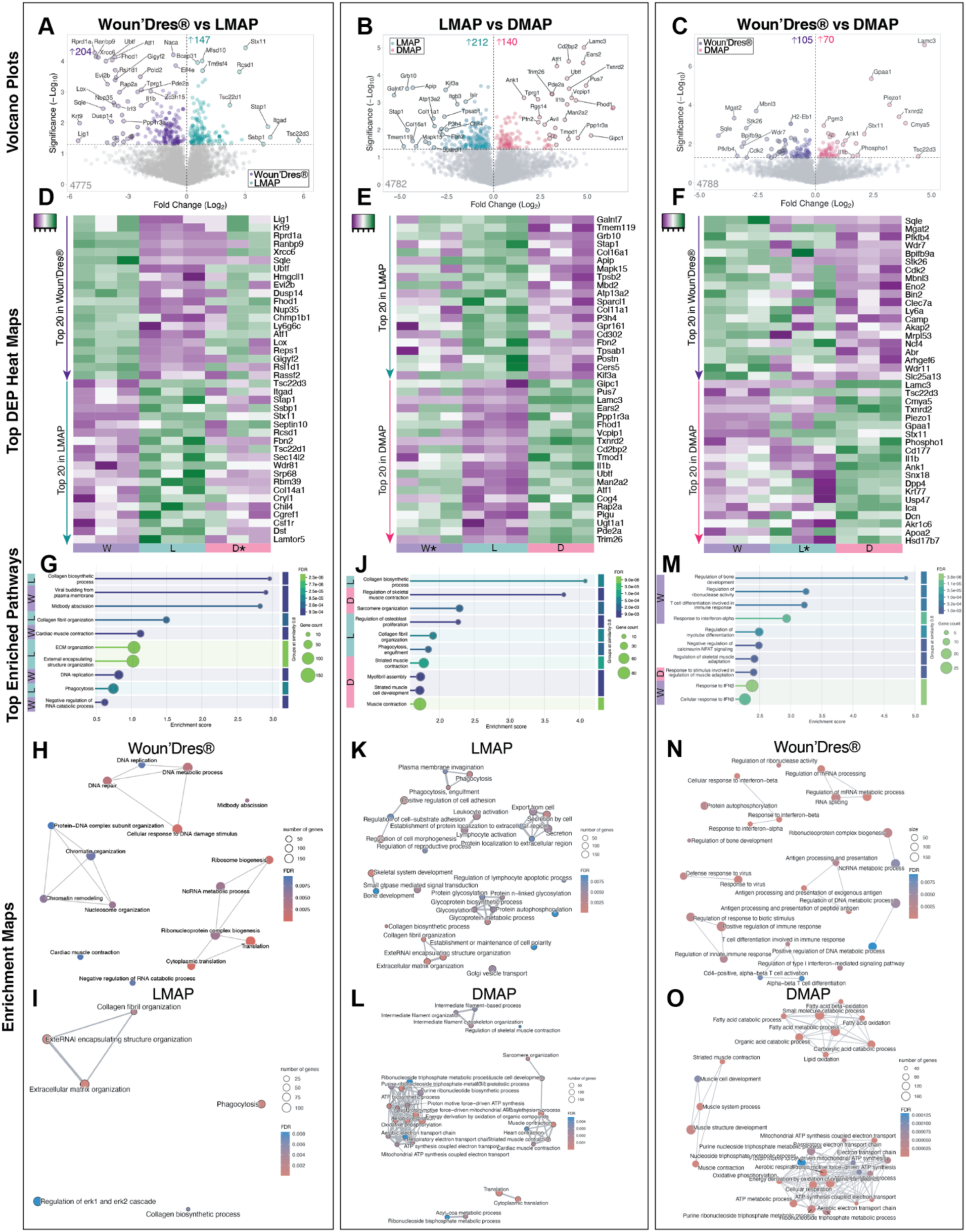
Proteomic analysis at day 14. **A-C)** Volcano plot showing the fold change (Log_2_) versus significance (-Log_10_ p-value) for proteins in comparisons **(A)** 1, **(B)** 2, and **(C)** 3. **D-F)** Heatmaps showing the expression levels of the 20 most upregulated proteins for each treatmentcomparing **(D)** Woun’Dres® vs LMAP, **(E)** LMAP vs DMAP, **(F)** Woun’Dres® vs DMAP. Proteins are ordered in decreasing order. **G)** Gene ontology enrichment analysis comparing biological processes between Woun’Dres® and LMAP, highlighting 10 of the most significantly enriched pathways. **H-I)** Enrichment GO pathways comparing **(H)** Woun’Dres® and **(I)** LMAP are indicated by node size (representing the number of genes) and color (indicating false discovery rate – FDR). **J)** Gene ontology enrichment analysis comparing biological processes between LMAP and DMAP, highlighting 10 of the most significantly enriched pathways. **K-L)** Enrichment GO pathways comparing **(K)** LMAP and **(L)** DMAP are indicated by node size (representing the number of genes) and color (indicating false discovery rate - FDR). **M)** Gene ontology enrichment analysis comparing biological processes between Woun’Dres® and DMAP, highlighting 10 of the most significantly enriched pathways. **N-O)** Enrichment GO pathways comparing **(N)** Woun’Dres® and **(O)** DMAP are indicated by node size (representing the number of genes) and color (indicating false discovery rate - FDR).

Notably, on Day 14, the DEPs showed reduced representation of proteins from the same families compared to the earlier time point, indicating a shift toward greater protein family diversity. Collagen was the most prominent protein family observed, with a consistent upregulation in LMAP within both Comparisons 1 (Col14a1) and 2 (Col16a1, Col11a1) **(Figure 4D-E**, **Supplemental Figure 6A)**, suggesting the upregulation of ECM deposition by LMAP. LMAP treatment also resulted in Fbn2 upregulation across the board **(Figure 4D-E**, **Supplemental Figure 6B)**, a protein that promotes microfibril formation during elastic fiber assembly^[55]^. Taken together, these results show that LMAP treated wounds have a high expression of proteins essential to granulation tissue deposition and the ECM.

Lamc3, a member of the Laminin family, was within the top 3 DEPs for DMAP in both Comparisons 2 and 3 **(Figure 4E-F**, **Supplemental Figure 6C).** As another component of the ECM, laminin is mostly found within the basement membrane, which is a major component of the dermal-epidermal junction (DEJ)^[56]^. Proper reconstitution of the DEJ is required for regenerative wound healing, including the DEJ undulating pattern^[57]^. This boundary has been highlighted as a signaling niche for keratinocytes residing in the epidermis, where it serves differing roles, including facilitating cell proliferation and differentiation^[58]^. While LMAP treatment more consistently upregulated ECM proteins, DMAP treatment instead resulted in the upregulation of an ECM protein known for its role related to hair follicles and keratinocytes, Lamc3^[59]^.

Tsc22d3 was the most DEP for LMAP and the second most DEP DMAP in Comparisons 1 and 3, respectively **(Figure 4D,F**, **Supplemental Figure 6D)**. This protein, also known as GILZ, has been recognized for its anti-inflammatory properties, partly through its ability to modulate the NF-κB pathway^[60]^. Contrary, DMAP also upregulated the inflammatory cytokine protein IL1*β* **(Figure 4E-F**, **Supplemental Figure 6E)**, with prolonged expression being associated with negative wound healing results^[61]^. This balance between pro-and anti-inflammatory nature has been previously observed for DMAP implants^[62]^.

Taken together the DEP analysis shows that LMAP supports protein upregulation associated with an anti-inflammatory environment and pro-ECM depositing environment, Woun’Dres® supports protein downregulation associated with an anti-inflammatory environment, and DMAP promotes a mixture of pro-and-anti-inflammatory environments with an upregulation of regenerative laminin.

### 2.7 Day 14 comparative Gene Ontology analysis highlights biomaterial dependent mechanisms

GO analysis of this phase yielded 23, 74, and 100 differentially expressed GO biological processes for Comparisons 1, 2, and 3, respectively **(Supplemental Data 7-9).** Of the biological processes highlighted by our comparative GO analyses on Day 14, LMAP had a consistent enrichment against both Woun’Dres® and DMAP of those related to ECM organization (‘Collagen fibril organization’, ‘Collagen biosynthetic process’) (**Figure 4G,I-K**, **Supplemental Figure 6G-H)** which were grouped into Regulation of Collagen in the Comparison 2 enrichment tree **(Supplemental Figure 6H)**. Woun’Dres® had terms related to Cellular DNA repair (‘DNA replication’, ‘DNA repair’, ‘DNA metabolic process’) against LMAP, indicating the distinct healing responses elicited by different biomaterial treatments **(Figure 4G-H**, **Supplemental Figure 6F)**. Instead in Comparison 3, Woun’Dres® enriched a variety of immune response pathways (‘T cell differentiation’, ‘Response to IFN*β*’), especially those related to viral response, when compared against DMAP **(Figure 4M-N**, **Supplemental Figure 6J).** Indicating the immune compartment as a strong enrichment difference between Woun’Dres® and DMAP on Day 14, despite the two biomaterials sharing similarities in this same category on Day 4.

DMAP treatment resulted in the enrichment of terms associated with muscle on Day 14 (‘Muscle contraction’, ‘Muscle system process’, ‘Muscle cell development’) **(Figure 6J, L-M, O**, **Supplemental Figure 6I, K)**. While not traditionally associated with dermal wound healing processes, we found this to be an interesting expression pattern conserved in both comparisons with DMAP. In Comparison 2, there were 11 terms repeated within the muscle-related family on Day 14, with 91% of them upregulated in DMAP alone and 9% upregulated in both treatment groups **(Supplemental Data 8)**. Muscle-related terms are not to be confused with wound contraction, which is mediated by myofibroblasts through the expression of alpha-smooth muscle actin (α-SMA). Excessive upregulation of this biological process has been associated with negative scarring outcomes^[63]^. This difference in muscle contraction and myofibroblast wound contraction was confirmed by the lack of the α-SMA protein (ACTA2) in the proteomes on Days 14 and 21 **(Supplementary Data 10-11)**.

### 2.8 At day 14 LMAP promotes upregulation of ECM- and collagen-related terms

The ECM is a crucial aspect of wound healing, with roles ranging from providing cellular support and influencing cell behavior to regulating the bioavailability of growth factors^[54a]^. Given our interest in the temporal aspect of wound healing, we revisited the Matrisome analysis technique with the Day 14 proteome. As wound healing progressed, ‘core matrisome’ proteins comprised 53 to 67% of total matrisome protein expression on Day 14 **(Figure 5A)**, compared to 23 to 29% on Day 4 **(Figure 3A)**. LMAP treatment significantly increased ‘core matrisome’ proteins compared to the other biomaterial treatments **(Figure 5A)**. Of the core matrisome categories, LMAP specifically had the greatest prevalence of collagens and ECM glycoproteins compared to the other wound treatments **(Figure 5B).**

**Figure 5.**
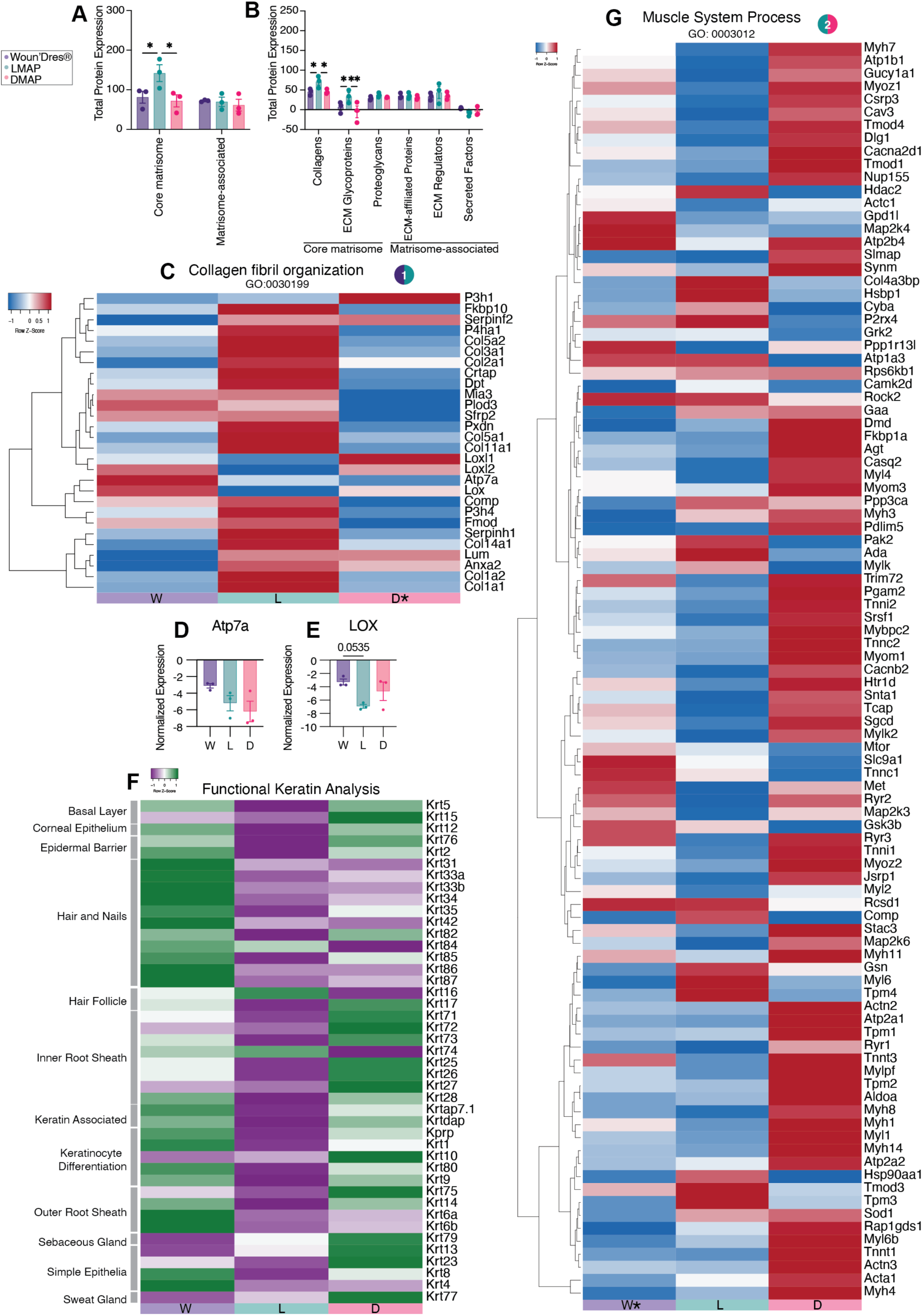
Detailed proteomic analysis of wound healing at day 14. **A)** Total protein expression in Core Matrisome and Matrisome-Associated Categories. Bar plots show total protein expression levels in the core matrisome and matrisome-associated proteins across the three experimental groups. Two-way ANOVA was completed with Tukey’s multiple comparisons test (simple effects within rows). Significant differences are indicated (*p < 0.05). **B)** Protein subcategories in Matrisome analysis. Detailed breakdown of protein expression within subcategories of Matrisome analysis, including collagens, ECM glycoproteins, proteoglycans, ECM-affiliated proteins, ECM regulators, and secreted factors. Two-way ANOVA was completed with Tukey’s multiple comparisons test (simple effects within rows). Significant differences between groups are marked (*p < 0.05). **C)** Collagen fibril organization (GO:0030199) Heatmap showing differential protein expression related to collagen fibril organization. Proteins are hierarchically clustered, with red indicating upregulation and blue indicating downregulation across the experimental group**s. D-E).** Normalized expression of key proteins. Bar plots display normalized expression levels of specific proteins involved in collagen organization**: (D)** Atp7a and **(E)** LOX. One-way ANOVA with Tukey’s multiple comparisons test was performed. **F)** Heatmap illustrating keratin protein expression associated with various skin structures and functions, such as the basal layer, corneal epithelium, epidermal barrier, hair follicles, and sebaceous glands. Expression levels are normalized and color-coded (green: low expression; purple: high expression). **G)** Muscle system process (GO:0003012) heatmap showing differential protein expression related to muscle process. Proteins are hierarchically clustered, with red indicating upregulation and blue indicating downregulation across the experimental groups.

We next explored the GO-identified term ‘Collagen fibril organization’ using a heatmap to analyze specific expression of genes within the term **(Figure 5C)**. LMAP had an upregulation of many collagen proteins, including Col141a1, Col1a1, Col1a2, Col2a1, Col5a2, Col3a1, Col5a1, Col11a1. Col1a1 and Col1a2 together represent Collagen I^[64]^, an essential component of the skin that maintains skin structure and integrity and accounts for up to 80% of the total collagen present in skin^[65]^. Woun’Dres® had an apparent upregulation of Atp7a and LOX **(Figure 5C)**. When the normalized expression of these proteins was plotted, Woun’Dres® had a trend for increased expression for Atp7a, though not significantly significant **(Figure 5D).** LOX had a similar increased expression trend in Woun’Dres® **(Figure 5E)**, with a P-value of 0.0023 when Woun’Dres® and LMAP were compared directly with a t-test. Lysyl oxidase (LOX) is part of a group of enzymes that aids in ECM stabilization through the catalysis of collagen and elastin crosslinking^[66]^. Excessive crosslinking of these ECM proteins results in scar formation^[67]^. The ATP7A protein is linked closely to the LOX family as it serves as a copper transporter required for proper LOX activity^[68]^. While LOX is an important part of ECM deposition and organization, it requires controlled expression. It has been previously shown that topical application of a small molecule inhibitor against LOX in a porcine excisional wound healing model reduces the appearance of scars^[69]^. These results further support the scarred versus ECM regenerated phenotypes seen in Day 21 histology and support LOX downregulation as a potential approach to improving tissue quality after injury.

### 2.9 At day 14 DMAP upregulates keratinocyte proteins and enhanced hair follicle formation

DMAP treatment pushed the wound environment towards the expression of proteins related to more developed hair follicles, including inner root sheath (IRS) keratins **(Figure 5F)**. During hair follicle formation, the outer root sheath is formed by epithelial cells proliferating downwards towards the dermis followed by the differentiation of the IRS into its various layers^[70]^. Of the two hair follicle keratins, DMAP and LMAP had opposing expression patterns. The expression of these keratins is known to be induced in skin after acute injuries and maintained until barrier function is restored, earning them the title of “stress keratins”^[71]^. Krt16 expression was upregulated in LMAP **(Figure 5F)** and is associated closely with keratinocyte migration at the wound edge^[72]^. Krt17 expression was upregulated in DMAP **(Figure 5F)** and is known to promote keratinocyte proliferation and migration as well^[73]^. Notably, the expression of this keratin is linked to hair follicle development during both embryogenesis^[74]^ and wound-induced hair follicle neogenesis^[75]^. Despite an earlier downregulation of keratin-related terms, DMAP treated wounds exhibited a strong jump in keratin proteins by Day 14, indicating Keratin expression as a tightly regulated wound response with DMAP treatment.

### 2.10 Day 14 shows an unexpected enrichment of muscle-related pathways in DMAP

With the surprising presence of muscle-related terms in DMAP, we used a heatmap of ‘Muscle system process’ to explore this pathway **(Figure 5G).** On Day 14, we found that DMAP had an upregulation of proteins involved in muscle contraction^[16b, 76]^(Tnnc, Tnni2, Myl4, Myh7, Mylpf and Mybpc2), muscle structure^[77]^ (Myom1, Synm, Dmd), and calcium handling^[78]^ (Casq2, Cacna2d1, Cacnb2, Atp2b4) **(Figure 5G)**. Additionally, one upregulated protein was highlighted with potential connections to wound healing: Trim72. Trim72, also known as MG53, is a muscle-enriched member of the TRIM family of E3 ubiquitin ligase proteins^[79]^. While highlighted as an essential component of the membrane repair machinery in muscle cells, exogenous delivery of Trim72 has been shown to promote therapeutic cell membrane repair in cell types beyond muscle cells^[80]^. Mice without Trim72 were shown to have baseline abnormal skin structure, with increased collagen content and decreased hair follicle density^[81]^. When a hydrogel was used to deliver exogenous Trim72 to the skin wounds of wild type mice, improved wound healing and visual scar reduction was observed^[81]^. Highlighting Trim72 as an interesting facilitator of rapid repair and reduced scar formation, beyond that of muscle related processes.

### 2.11 Day 21 comparative Gene Ontology analysis highlights biomaterial dependent mechanisms

As the healing process progressed towards resolution, differences between MAP and Woun’Dres® were apparent; comparisons against LMAP and DMAP yielded 501 and 498 total DEPs **(Figure 6A,C)**. This contrasts the similarities between the two MAP treatments, with only 158 total significant DEP proteins **(Figure 6B)**. Similar to Day 14, there were not many protein families that were repeatedly upregulated in the top 20 DEP heatmaps **(Figure 6D-F).** One superfamily which was consistently upregulated by DMAP treatment was the Solute Carrier (Slc) family, including Slc25a42 in both Comparison 2 and 3 **(Figure 6E-F**, **Supplemental Figure 7A).** This protein assists in the transportation of coenzyme A (CoA) into mitochondria for various metabolic processes^[82]^. We observed many (17) repeated proteins upregulated across comparisons. Of these, we were particularly interested in the upregulation of Lamc3 by both LMAP and DMAP in comparisons against Woun’Dres® **(Figure 6D,F**, **Supplemental Figure 7B)**. Originally Lamc3 was highly expressed in DMAP alone on Day 14 **(Figure 4E-F)**. This shift in expression pattern corroborates the similarity in MAP profiles at this later time point.

**Figure 6.**
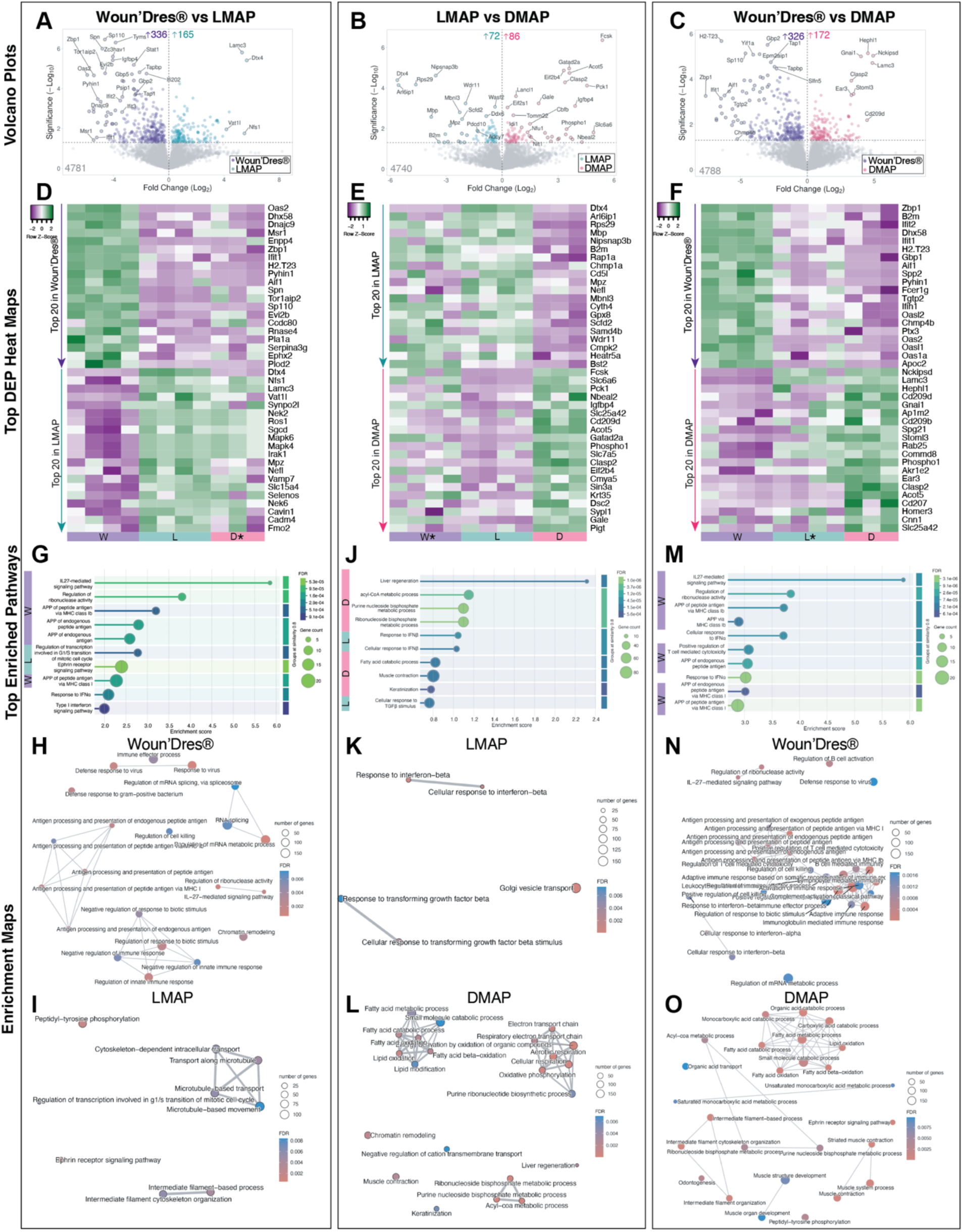
Proteomic analysis at day 21. **A-C)** Volcano plot showing the fold change (Log_2_) versus significance (-Log_10_ p-value) for proteins in comparisons **(A)** 1, **(B)** 2, and **(C)** 3. **D-F)** Heatmaps show the expression levels of the 20 most upregulated proteins comparing **(D)** Woun’Dres® vs LMAP, **(E)** LMAP vs DMAP, and **(F)** Woun’Dres® vs DMAP. Proteins are ordered in decreasing order. **G)** Gene ontology enrichment analysis comparing biological processes between Woun’Dres® and LMAP, highlighting 10 of the most significantly enriched pathways. **H-I)** Enrichment GO pathways comparing **(H)** Woun’Dres® and **(I)** LMAP are indicated by node size (representing the number of genes) and color (indicating false discovery rate - FDR). **J)** Gene ontology enrichment analysis comparing biological processes between LMAP and DMAP, highlighting 10 of the most significantly enriched pathways. **K-L)** Enrichment GO pathways comparing **(K)** LMAP and **(L)** DMAP are indicated by node size (representing the number of genes) and color (indicating false discovery rate - FDR). **M)** Gene ontology enrichment analysis comparing biological processes between Woun’Dres® and DMAP, highlighting 10 of the most significantly enriched pathways. **N-O)** Enrichment GO pathways comparing **(N)** Woun’Dres® and **O)** DMAP are indicated by node size (representing the number of genes) and color (indicating false discovery rate - FDR).

Dtx4 was the top upregulated protein in LMAP against both other biomaterial treatments **(Figure 6D-E**, **Supplemental Figure 7C).** As mentioned earlier, Dtx4 was uniquely expressed in MAP treatments at later times points (Day 14 DMAP, Day 21 LMAP) and is linked to ubiquitin ligase activity. We observed the repeated upregulation of various immune proteins by Woun’Dres® in Comparisons 1 and 3 **(Figure 6D,F).** These included both Ifit1 **(Supplemental Figure 7D)**, an antiviral protein induced by interferons^[83]^, and H2-T23 **(Supplemental Figure 7E)**, an MHC class 1b protein^[84]^, suggesting that the immune response to Woun’Dres® was not resolved by Day 21. Interestingly, DMAP treatment upregulated an enzyme linked to fatty acid metabolism, Acot5 **(Figure 6E-F**, **Supplemental Figure 7F).** The Day 21 profile of MAP treated wounds boasted an array of protein upregulations whereas Woun’Dres® wounds upregulated proteins associated with the immune response, akin to its earlier profiles.

### 2.12 MAP enriches a diversity of biological processes on Day 21

A total of 31, 27, and 80 differentially expressed GO biological processes were identified for Comparisons 1, 2, and 3, respectively **(Supplemental Data 10-12)**. Woun’Dres® maintained the inflammatory environment seen on Day 4 through Day 21, with upregulation of terms such as ‘Antigen processing and presentation (APP) of peptide antigen’, and ‘Immune effector process’ which were repeated in both Comparisons 1 and 3 **(Figure 6G-H, M-N**, **Supplemental Figure 7G, K)**. Interestingly, ‘IL-27 mediated signaling pathway’ was the top enriched term for Woun’Dres® for both comparisons **(Figure 6G,M).** Prolonged inflammation is a hallmark of impaired wound healing, frequently observed in conditions such as diabetes, and has been correlated with increased scarring^[85]^. While Woun’Dres® had a consistent immune upregulated profile, both MAP treatments had diverse biological enrichments **(Figure 6G,I-M,O)**.

In Comparison 1, the LMAP proteome included upregulated terms related to the cytoskeleton and intermediate filaments (‘Intermediate filament cytoskeleton organization’, ‘Microtubule-based transport and movement’) **(Figure 6G,I**, **Supplemental Figure 7H).** The cytoskeleton is the framework of the cell which is comprised of three systems: actin microfilaments, intermediate filaments, and microtubules^[86]^. Intermediate filaments are an extremely flexible component of the cytoskeleton, which benefits the cell during late stage wound healing by maintaining cell integrity and mechanical stability during tissue remodeling^[87]^. Microtubules are essential for intracellular transport and cell migration, allowing for signaling proteins and vesicles to traffic and contribute to coordinated repair. Interestingly, DMAP’s proteome had a similar intermediate filament enrichment scheme against Woun’Dres® **(Figure 6O**, **Supplemental Figure 7L)**, indicating that MAP treatment improves cell and mechanical integrity by this wound resolution time point, as was shown previously^[25b]^.

In accordance with the small number of DEPs and GO processes enriched between the two MAP conditions, Comparison 2 boasted the smallest enrichment scores **(Figure 6J).** In Comparison 2, the LMAP profile changed, with an enrichment of pathways important for ECM remodeling^[88]^ (‘Response to TGF*β*’) and the inflammatory response^[89]^ (‘Response to IFN*β*’) **(Figure 6J-K**, **Supplemental Figure 7I).** Within this same comparison, DMAP enriched lipid (‘Fatty acid catabolic process’, ‘Fatty acid oxidation’), muscle (‘Muscle contraction’), and keratin (‘Keratinization’) related pathways **(Figure 6J,L**, **Supplemental Figure 7J).** Keratinization occurs specifically in the stratum corneum and hair which correlates well with the increased presence of hair follicles in DMAP treated wounds on Day 21 **(Supplemental Figure 2)**. DMAP treatment similarly enriched these muscle and lipid processes when compared against Woun’Dres® **(Figure 6O**, **Supplemental Figure 7L).** Notably, none of the terms enriched by DMAP treatment on Day 21 **(Supplemental Data 11-12)** were directly associated with inflammation or the immune response within the wound bed proteome.

### 2.13 At day 21 MAP treatment results in improved collagen despite enriching differential pathways throughout healing

Inspired by the upregulation of ECM related terms seen in MAP compared to Woun’Dres®, we further explored the ECM composition during the remodeling time points of wound healing using the same Matrisome bioinformatics approach **(Figure 7A-B)**. The trend of core matrisome components becoming more prevalent seen on Day 14 **(Figure 5A)** continued through Day 21, with core matrisome comprising 72-92% of total matrisome proteins **(Figure 7A).** Interestingly, all treatments expressed similar total amounts of the core components **(Figure 7B)**, indicating that the treatment profiles became more similar at this later time point. ECM regulators were significantly upregulated on Day 21 in Woun’Dres® **(Figure 7B)**. This matches our observations that Woun’Dres® had similar proteome profiles on Day 4 and Day 21 – first observed with the immune biological processes and then observed with the ECM compartment.

**Figure 7.**
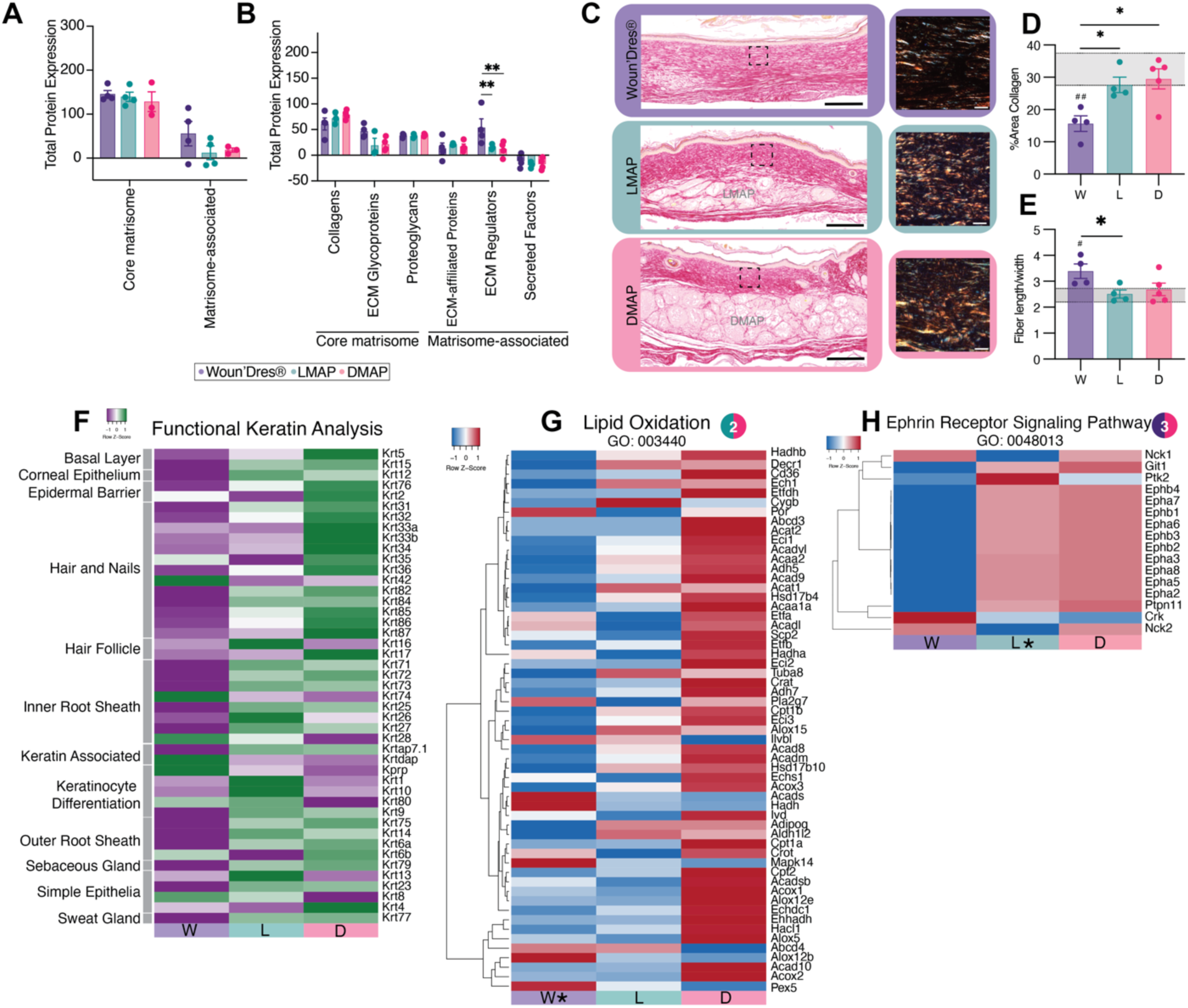
Detailed proteomic analysis of wound healing at day 21. **A)** Total protein expression in Core Matrisome and Matrisome-Associated Categories. Bar plots show total protein expression levels in the core matrisome and matrisome-associated proteins across three biomaterial treatment groups. Two-way ANOVA was completed with Tukey’s multiple comparisons test (simple effects within rows). Significant differences are indicated (*p < 0.05, **p < 0.01). **B)** Protein subcategories in Matrisome analysis. Detailed breakdown of protein expression within subcategories of matrisome analysis, including collagens, ECM glycoproteins, proteoglycans, ECM-affiliated proteins, ECM regulators, and secreted factors. Two-way ANOVA was completed with Tukey’s multiple comparisons test (simple effects within rows). Significant differences between groups are marked (*p < 0.05). **C)** Representative histological samples stained using Pricosirus red (PSR) imaged with brightfield (left) and polarized light (right) from Day 21 wounds. Gray text shows MAP location. Black box highlights region polarized light images were captured. Scale bar is 200 µm for brightfield and 20 µm for polarized light. **D-E)** Quantification of collagen features using PSR stained wounds. Grey bars represent wild-type skin results and the standard deviation. Hash-tag represents significance compared to wild-type. One-way ANOVA with Tukey’s multiple comparisons test was performed. Significant differences between groups are marked (*,#p < 0.05, ##P < 0.01). **F)** Heatmap illustrating keratin protein expression associated with various skin structures and functions, such as the basal layer, corneal epithelium, epidermal barrier, hair follicles, and sebaceous glands. Expression levels are normalized and color-coded (green: low expression; purple: high expression). **G-H)** Heatmap showing differential protein expression related to **(G)** Lipid oxidation (GO: 003440) and **(H)** Ephrin receptor signaling pathway (GO: 0048013). Proteins are hierarchically clustered, with red indicating upregulation and blue indicating downregulation across the experimental groups.

We sought to correlate the ECM proteomic findings with our histology data using Picrosirus red (PSR) staining, a technique for analyzing collagen quality that capitalizes on the birefringence of PSR-stained collagen when imaged under polarized light^[90]^. In MAP-treated wounds, we observed a more basket weave-like collagen organization that is characteristic of unwounded tissue versus tightly packed and parallel-oriented fibers in Woun’Dres® **(Figure 7C**, **Supplemental Figure 8A-C)**. When quantified **(Figure 7D)**, both LMAP and DMAP demonstrated comparable collagen area to that of normal skin, whereas Woun’Dres® collagen fibers were significantly more sparse compared to normal skin and both MAP treatments. Additionally, Woun’Dres® had a significantly increased collagen fiber length to width ratio compared to control **(Figure 7E).** LMAP and DMAP both had a lower collagen fiber length to width ratio than Woun’Dres®, closer to that of normal skin, although DMAP’s was not significant **(Figure 7E)**. This indicates a thicker, less scar like collagen fiber in MAP treated wounds. Despite LMAP having a more consistent enrichment of ECM related pathways throughout the wound healing time course, both MAP treatments ultimately resulted in similar improved collagen architecture.

### 2.14 DMAP is a top upregulator of Keratin related proteins on Day 21

Despite an earlier (Day 4) downregulation of keratinization-related terms, DMAP treated wounds were top producers of proteins found within this pathway on Day 21, as demonstrated by the identification of the ‘Keratinization’ pathway through GO analysis against LMAP **(Figure 6J,L)**. This pathway is downstream of ‘Epidermis development’ and ‘Keratinocyte differentiation’. On Day 21, DMAP’s keratin profile included those related to epidermal barrier, hair and nails, sebaceous glands, and outer root sheath **(Figure 7G)**. This shift towards the resolution of wound healing matches our observations of improved regenerative outcomes in DMAP-treated wounds. Notably, LMAP and DMAP maintained their differential upregulation of Krt16 and Krt17 (**Figure 7F)** seen on Day 14 **(Figure 5F).**

### 2.15 At Day 21 fatty acid metabolism is increased in DMAP

An underappreciated process in skin wound healing that was identified in the proteome of DMAP treated wounds is lipid modification on both Days 14 (**Figure 4O**) and 21 **(Figure 6L,O).** Lipids play important roles in the skin via barrier function and promoting recovery as well as regeneration^[91]^. Heatmap analysis of the ‘Lipid oxidation’ GO pathway showed an increased presence of proteins within this pathway in DMAP **(Figure 7H)**. Within this pathway, DMAP specifically upregulated Cyp4f18, ALOX5, and ALOX12E **(Figure 7G)**, all of which are important enzymes involved in fatty acid lipid metabolism with their products playing a role in inflammation modulation^[92]^. Additionally, ACAD, CPT1A, CPT1b, CPT2, ECHDC1, and CRAT were upregulated by DMAP **(Figure 7G)**. These proteins are linked with fatty acid oxidation and metabolism in ways that produce energy necessary to fuel cellular processes important to wound healing^[93]^. We highlighted the upregulation of Krt17 earlier in regard to its role in regeneration, but it has also been linked with increased lipid synthesis for promoting improved barrier reinstatement^[94]^.

### 2.16 Granular scaffolds promote the Ephrin pathway at Day 21

While LMAP and DMAP performed differently across the timepoints, there was one pathway clearly identified to be common during the matrix remodeling phase. On Day 21, the ‘Ephrin receptor signaling pathway’ was clearly upregulated in both MAP conditions but not Woun’Dres® **(Figure 7H)**. These receptors are part of the largest family of receptor tyrosine kinases and boast a unique bidirectional signaling between receptor and ligand^[95]^. Their downstream signaling pathways including Rho GTPases, MAP kinases, and PI3K have a variety of roles in wound healing, impacting cell adhesion, motility, proliferation and differentiation^[96]^. Ephrin receptors have been suggested to impact many cell types, such as cells resident to the skin including keratinocytes, Langerhans cells, dermal fibroblasts, and lymphocytes^[97]^. Interestingly, EphB signaling is implicated in loosening adhesion junctions between epidermal cells to facilitate re-epithelization, as well as in facilitating keratinocytes to release epithelial tension through disassembly of contractile stress fibers^[98]^. Overexpression of the ephrin pathway, however, has been associated with fibrotic responses in skin^[99]^, though this phenotype was not observed with MAP treatment. These literature reports leave much to explore in this MAP specific pathway.

## 3. Conclusion

Through this work, we presented the temporal responses to different biomaterial treatments which facilitate varying degrees of skin regeneration **(Figure 8)**. We investigated Woun’Dres®, LMAP, and DMAP across Days 4, 14, and 21 to capture a holistic view of the wound healing cascade using a comprehensive proteomic approach with analysis focused on matrisome content, functional keratin analysis, and enrichment of GO biological pathways.

**Figure 8.**
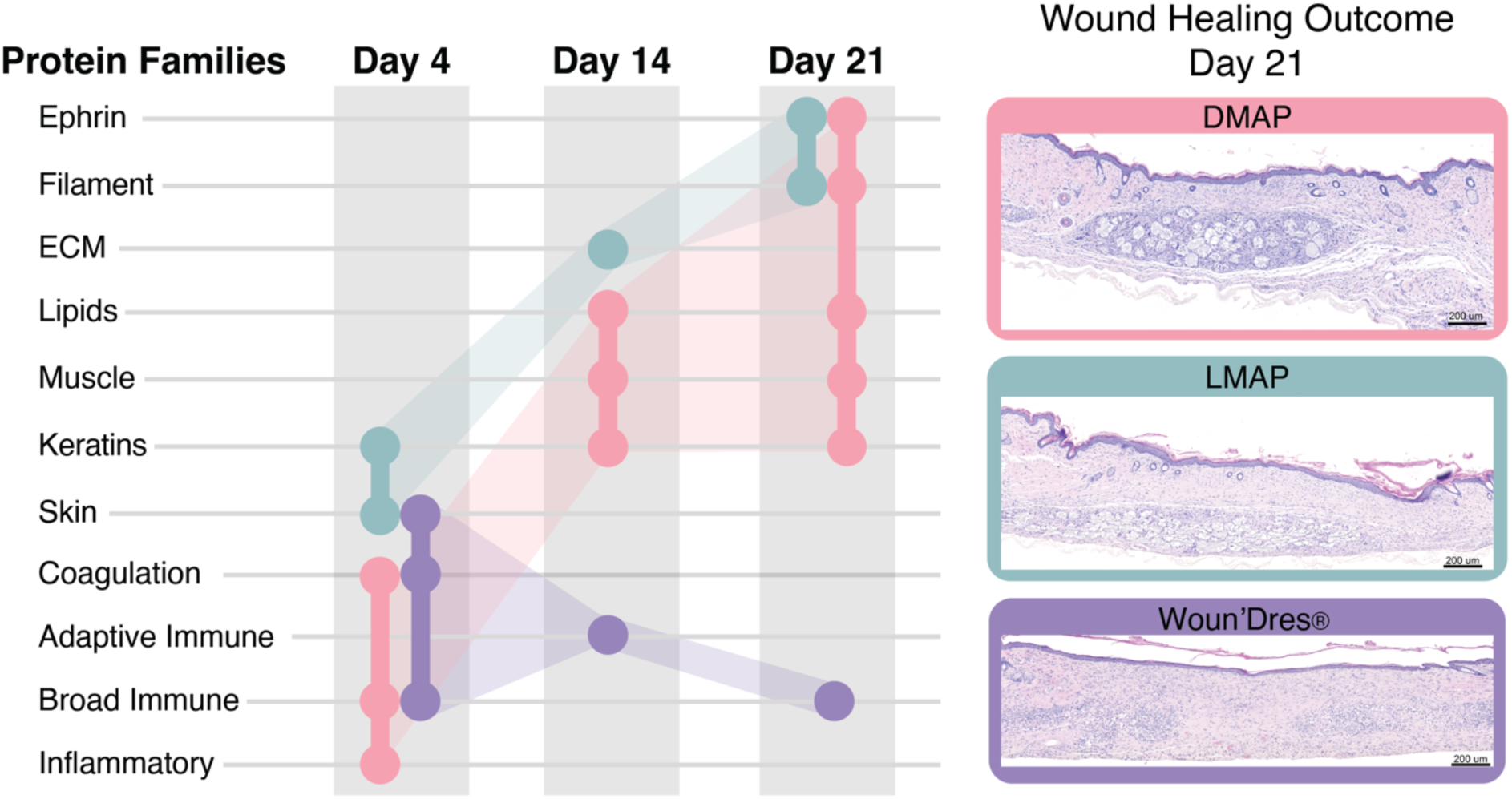
Temporal dynamics of protein families during wound healing and histological outcomes at day 21. The left panel shows the different protein families involved in wound healing (y-axis) across three time points: Day 4, Day 14, and Day 21 (x-axis). Colored lines represent distinct treatment groups: DMAP (pink), LMAP (teal), and Woun’Dres® (purple). The right panel presents representative histological images of wound sites at Day 21 for each treatment group (DMAP, LMAP, Woun’Dres®). Images are stained with hematoxylin and eosin (H&E) and scale bars represent 200 μm. The images illustrate differences in tissue architecture and healing quality among treatments.

We found that wounds separated by timepoint in PCAs, highlighting the importance of the temporal component of wound healing. We first investigated the biomaterial induced proteome at Day 4. DMAP and Woun’Dres® profiles were similar in the enrichment of both broad immune pathways as well as coagulation pathways. Given the difference in wound morphology at Day 21 and the vastly different biomaterial composition, this alignment of the two biomaterials was surprising. Although, their profiles did not completely overlap, as it was noted that DMAP in fact enriched the inflammatory response when compared to both Woun’Dres® and LMAP. Adaptive immunity was previously shown to be essential in DMAP mediated regeneration as regeneration was lost in a RAG1^-/-^ mouse model lacking all mature B and T cells^[25b]^. Yet the window in which the immune response to DMAP is most essential was not elucidated before this proteomic study. On the other hand, the early wound proteome of LMAP was sufficiently different from both Woun’Dres® and its D-AA counterpart, promoting more terms related to keratinocyte differentiation and protein synthesis. LMAP and Woun’Dres® both upregulated skin developmental pathways compared to DMAP, highlighting the consistent downregulation by DMAP. Since Day 4 is within the inflammation window, it seems that LMAP facilitates a quick transition away from inflammation towards the proliferation phase, whereas DMAP and Woun’Dres® promote immune engagement.

When the wounds were within the proliferative window, LMAP promoted a strong upregulation of ECM-related pathways, most notably with collagen organization and core matrisome components. This profile was maintained regardless of the biomaterial which LMAP was compared to. By Day 14, the proteomes of Woun’Dres® and DMAP treated wounds diverged. When these two treatments were compared, Woun’Dres® dominated in the enrichment of immune terms, especially those related to adaptive immunity, whereas DMAP interestingly enriched muscle and lipid related processes. Notably, functional keratin analysis highlighted DMAP as a top up regulator of proteins associated with more mature hair follicle phenotypes.

The inflammatory environment seen in Woun’Dres® prevailed on to Day 21, with the wound environment returning to enrichment of broad immune processes. DMAP again enriched muscle and lipid related processes whereas LMAP enriched TGFβ/ILβ against DMAP. Despite a Day 4 downregulation of keratinization-related terms, DMAP treated wounds were top producers of proteins found within this pathway again on Day 21. MAP treatment, regardless of crosslinking peptide chirality, enriched the Ephrin receptor pathway as well as intermediate filament related pathways.

Overall, we highlighted important biological process and protein expression differences in dermal full-thickness wounds treated with three biomaterials: two similar in composition other than the chirality of amino acids within the crosslinking peptide and one which differed completely in material composition. Both MAP treatments resulted in improved regenerative outcomes which were reached by vastly different pathways over the course of healing. The LMAP proteome initially enriched epidermal differentiation pathways but moved towards ECM enrichment when the wound was surveyed on Days 14 and 21. The DMAP proteome began with local inflammation and immune activation but then moved towards processes related to wound closure and regeneration, with a stark switch in keratin protein profiles between Days 4 to later time points. Woun’Dres® resulted in a scar like phenotype which was instigated in part through a sustained inflammatory response starting on Day 4 and carried through to the resolution phase. A greater understanding of the pathways regulated by regenerative biomaterials provides critical information to not only understanding MAP’s regenerative mechanisms but also learning more about how chirality and biomaterials can be engineered for regenerative medicine.

## 4. Materials and Methods

### 4.1 Microparticle generation and purification

Microfluidic devices and microgels were produced as previously described^[25b]^. Briefly, 4-arm Polyethylene Glycol-Vinylsulfone (PEG-VS, 20kDa, JenKem) was dissolved in 0.3 M triethyloamine (Sigma) pH 8.8 and pre-reacted with K-peptide (Ac-FKGGERCG-NH2, GenScript), Q-peptide (Ac-NQEQVSPLGGERCG-NH2, GenScript) and RGD (Ac-RGDSPGERCG-NH2, GenScript) for at least one hour at 37 °C. Then, the pH was adjusted to 4.5-5.5 to slow down the on-chip gelation and prevent any clogging at the flow-focusing region. The final precursor concentration was 10% (w/v) 4-arm PEG-VS with 500 μM K-peptide, 500 μM Q-peptide, and 1000 μM RGD. The cross-linker solution was prepared by dissolving the di-thiol matrix metalloproteinase sensitive peptide L-MMP(Ac-GCRDGPQGIWGQDRCG-NH2, GenScript) or D-MMP (Ac-GCRDGPQ_D_GI_D_W_D_GQDRCG-NH2, GenScript) in distilled water at 16 mM for 4-arm LMAP or 16.4 mM for 4-arm DMAP and reacted with 10 μM Alexa-Fluor 647-maleimide (Thermo Fisher) for 5 minutes. These solutions were filtered through a 0.22 μm sterile filter, mixed 1:1, and loaded into a 1 ml syringe. The pinching oil phase was a heavy mineral oil supplemented with 1% v/v Span-80 (Sigma Aldrich). Downstream of the segmentation region, a second oil inlet with a high concentration of Span-80 (5% v/v) and Triethylamine (3% v/v, Sigma Aldrich) was added and mixed to the flowing droplet emulsion to raise pH and initiate cross-linking. The final HMP was collected in a conical tube and allowed to complete gelation at room temperature overnight. After gelation, the HMP and oil mixture was centrifuged to remove the oil phase top layer. The HMPs were then washed with 2% Pluronic F127 (Sigma Aldrich) in HEPES buffer (pH 8.3, 0.3 M) to assist with removal of the surfactant. Follow up washes were performed with HEPES buffer (pH 8.3, 0.3M, 1% Antibiotic-antibiotic (AA)) and centrifugation at max speed. The purified microspheres were tested to ensure no endotoxin (Pierce) and then stored in HEPES buffer (pH 8.3, 0.3M, 1% AA, 10 mM CaCl_2_) at 4 °C.

### 4.2 Material Property Characterization

Material properties were tested using a Rheometer (Anton Paar). Amplitude sweeps were first performed to identify the viscoelastic region. The identified percent strain (4% for MAP scaffolds, 6% for Woun’Dres®) was then used for all frequency sweeps for storage modulus.

### 4.3 In vivo wound healing study

Mixed sex SKH-1 Elite mice (8-12 weeks old) were purchased from Charles River and housed in a centralized animal facility at Duke University. The excisional splinted wound protocol is an established methodology previously detailed by us and others^[12a, 25b, 100]^. All animal work was performed under an IACUC approved protocol. Briefly, the mice were anesthetized with 4% isoflurane (1.5–2% isoflurane for maintenance) and placed on a heating pad. Buprenorphine SR (one dosage of 0.5 mg/kg of mouse weight, 0.5 mg/mL concentration) was injected subcutaneously. The dorsal surface of mice was prepped with alternating washes of iodine and 70% isopropyl alcohol. Using a sterile 5 mm biopsy punch, two through and through punches were made, creating four clean, well-defined wounds along the middle of the animal’s back. Adherent PDMS (aPDMS) ring splints with a 7-mm wide internal window were adhered around the wound to prevent contraction and allow for healing through re-epithelialization and granulation.^[100b]^ The HMPs were dried via centrifugation and removal of supernatant the day of surgery. They were then mixed with final concentrations of 4 U/ml of thrombin (200 U/mL in 200 mM Tris-HCl, 150 mM NaCl, 20 mM CaCl_2_) and 10 U/ml of Factor XIII (250 U/mL) and immediately injected to the wounds (10 µL/wound) for in situ gelation of the MAP scaffolds. Woun’Dres® was as manufacturer recommended (10 µL/wound). Of the four wounds on the mouse, they received a random combination of DMAP, LMAP or Woun’Dres® collagen hydrogel. After a 30-minute gelation time, Tegaderm dressings were adhered to both seal the splints and cover the wounds to prevent the wounds from drying out and to protect them from scratching. Animals were housed individually in cages with proper enrichment. Animals were checked on daily to ensure proper weight maintenance, as well as splint and bandage adherence. On days 4, 14 and 21, animals were euthanized, and wounded tissues were cut out for histology or proteomic profile.

### 4.4 Sample collection

Mice were euthanized and the back skin was carefully removed. 1.5-mm biopsy punch samples were collected from the center of the wounded tissue and stored in 1.5 mL low binding tubes. For paraffin embedding of wounds, 8-mm biopsy punch samples around the wounding site were collected from euthanized mice on Day 21 and fixed with 4% paraformaldehyde overnight at 4 °C before paraffin embedding. Paraffin embedded wounds were cut into 5-μm sections for histology staining.

### 4.5 Histological tissue staining

For all histological stains, sections were first deparaffinized and rehydrated using decreasing concentrations of xylene and ethanol. Hematoxylin and Eosin (H&E) staining proceeded with staining in Mayer hematoxylin solution (EMS) for 15 minutes followed by the blue-ing step with warm tap water for 15 minutes. Slides were then washed with D.I. water and 95% ethanol before staining in alcoholic eosin Y counterstain (EMS) for 40 seconds before mounting steps. PSR staining proceeded with staining in Picrosirius red for 60 minutes. Slides were then washed with two changes of acidified water before mounting steps. After staining was completed for all staining types, slides were dehydrated and cleared in increasing concentrations of ethanol and xylene. They were then mounted in Permount (Fisher Scientific). Brightfield images were taken using a Zeiss Axio Scan at 20X. Polarized light images were taken using a Zeiss microscope with a polarized light filter with three images taken within the wound region at 40X for each section.

### 4.6 Histological analysis

All tissue analysis was performed in a blinded manner. Histological analysis of dermal appendages was performed with the help of a dermapathologist. The number of neogenic hair follicles within the wound bed was counted then divided by overall wound length for standardization. The afollicular percentage was calculated by finding the greatest distance between two dermal follicular appendages and then dividing it by the overall wound length to find the percent of the wound bed without hair follicles. Analysis of the PSR was done using ImageJ as described previously^[23]^. Briefly, images were opened in ImageJ and the color balance was corrected for each channel using the auto function. The channels were then split, and the red channel was used for analysis. Auto threshold was run on a population of images from the experiment to identify the ideal threshold (Li Threshold on Auto). The images were then thresholded and two areas (approximately 50 µm x 50 µm squares, using the Polygon tool to help avoid any biomaterial or dermal features) were selected. The ‘Analyze Particles’ tool was used with a 0.5 to infinity size limit to reduce noise. Results were then averaged across the 3 images with two areas within each image analyzed for %Area, Major (Length), and Minor (Width).

### 4.7 Protein digestion and LC-MS/MS analysis

Protein digestion and LC-MS/MS analysis was completed in collaboration with the Muddiman Lab at North Carolina State University.^[101]^ In brief, the 1.5 mm samples were subject to homogenization with 200 µL 50 mM ABC with 1% SDC, with 2.8 mm beads, 7 mins at 1500xg speed. Then, protein amount was measured by A280 and a volume equivalent to 40 µg was subsequently taken through Filter Aided Sample Preparation (FASP) with tryptic digestion for bottom-up proteomics. After completion of FASP, the samples were lyophilized and stored at −20°C until needed. The digested samples were analyzed using an EASY nanoLC-1200 interfaced to a Thermo Scientific Exploris-480 with an EASY-SprayTM mass spectrometer (Elite Version, Thermo Scientific, San Jose, USA) coupled with an EASY-nLC system (Thermo Scientific, USA) by a nano electrospray ion source. Then, Proteome Discoverer or PD (Thermo Scientific, San Jose, CA) 2.4 was used for database searching of the experimental nanoLC-MS/MS data against reviewed uniport mouse skin database.

### 4.8 Proteomic analysis

Data processing: Data Transformation, normalization, imputation, fold change and statistical test were performed following the guidelines described by Aguilan et al^[27]^. Filtering out of highly variant biological replicas was done using hierarchal clustering with average linkage and Euclidean distance. Outliers were identified as samples above a threshold defined by 1.5 times the Interquartile Range (IQR) above the third quartile of the dendrogram heights^[102]^. Classification, functional enrichment, gene ontology analysis, protein interaction analysis, and GO hierarchical analysis were performed using STRING (v12)^[36]^ pairing with clusterProfiler^[103]^.

Plotting: Volcano plots were made using VolcanoNoseR^[104]^. All heatmaps were made using the expression function on Heatmapper.ca. All heatmap data is scaled to row (protein). Heatmaps from GO pathways used GO pathways within the top 35% of enriched pathways. Pooled heatmaps were created by taking the average expression of a protein for each replicate. Matrisome analysis was completed using MatrisomeAnalyzer ^[39]^ and the outputs were plotted in Prism for visualization. PCA plots were made using the PCA function in Prism (version 10.4.1). Venns diagram plots were made using InteractiVenn^[105]^.

### 4.9 Biological replicates and statistics

For all *in vivo* studies, each data point represents a biological replicate of a wound from a different mouse treated with the same biomaterial. For all experiments, significance is indicated by \**P* < 0.05, \*\**P* < 0.01, \*\*\**P* < 0.001, \*\*\*\**P* < 0.0001. All statistical analyses were performed using Graphpad Prism software. All data is represented with mean +/- SEM. Specific statistical analyses are listed in the according figure captions.

## Acknowledgements

This research was supported by the National Institutes of Health under grants R01AI152568 and R01GM087964. Mass spectrometry measurements were conducted at the Molecular Education, Technology and Research Innovation Center (METRIC) at North Carolina State University. Thank you to Dr. Michelle Schneider, Assistant Professor of Pathology at Duke University, for her expertise and advice in histological analysis of tissue sections. Thanks to Amy Kim for her help in acquiring the polarized light images of the wounded tissue. We would like to thank Dr. Cameron Miller and Dr. Rachel Myers from the Bioinformatics and Clinical Analytics Team (BioCAT) at Duke for their support and guidance in the proteomic analysis. We would also like to thank Dr. Eric Monson from the Center for Data and Visualization Sciences at Duke for his support in visualizing the conclusion figure. Lastly, we would like to thank Dr. Dianne Cruz and the White-McGarrah Lab at Duke University for providing access to their AxioScan facility.

## Supplemental figures

**Supplemental Figure 1.**
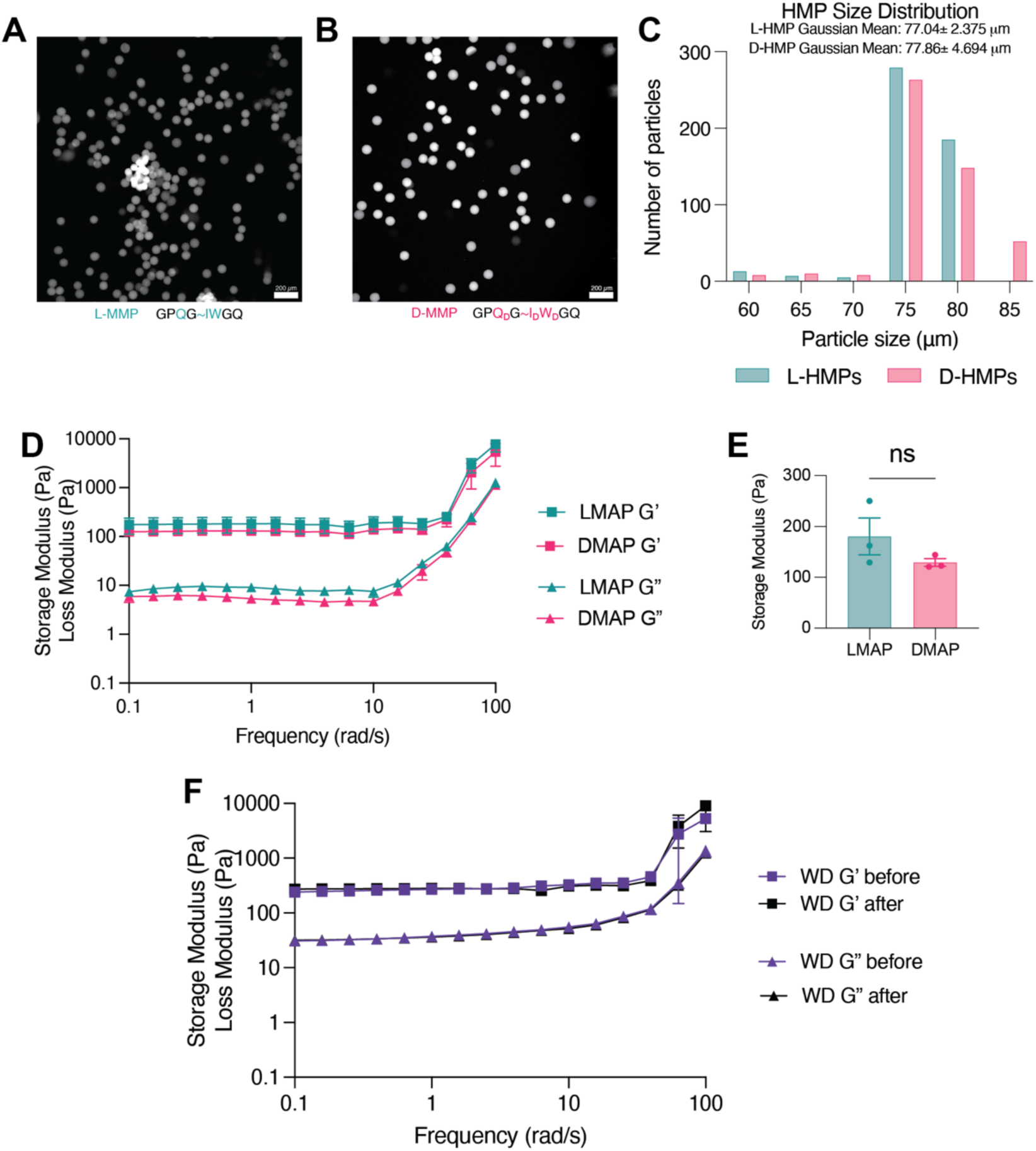
Characterization of size and mechanical properties of LMAP and DMAP HMPs and Woun’Dres® Hydrogel. **A-B)** Representative fluorescence microscopy images of (A, teal) LMAP and (B, pink) DMAP hydrogel microparticles (HMPs), showing particle morphology and distribution. Scale bars: 200 μm. **C)** Size distribution of L-HMPs (teal) and D-HMPs (pink), with Gaussian means indicated for each group. **D)** Frequency sweep rheology showing storage modulus (G’) and loss modulus (G’’) for LMAP and DMAP hydrogels, indicating similar viscoelastic properties. **E)** Quantification of storage modulus (G’) for LMAP and DMAP hydrogels; no significant difference observed using an unpaired t test (ns: not significant). **F)** Frequency sweep rheology of Woun’Dres® hydrogel before and after 30 minutes at 37°C, demonstrating consistent viscoelastic properties.

**Supplemental Figure 2.**
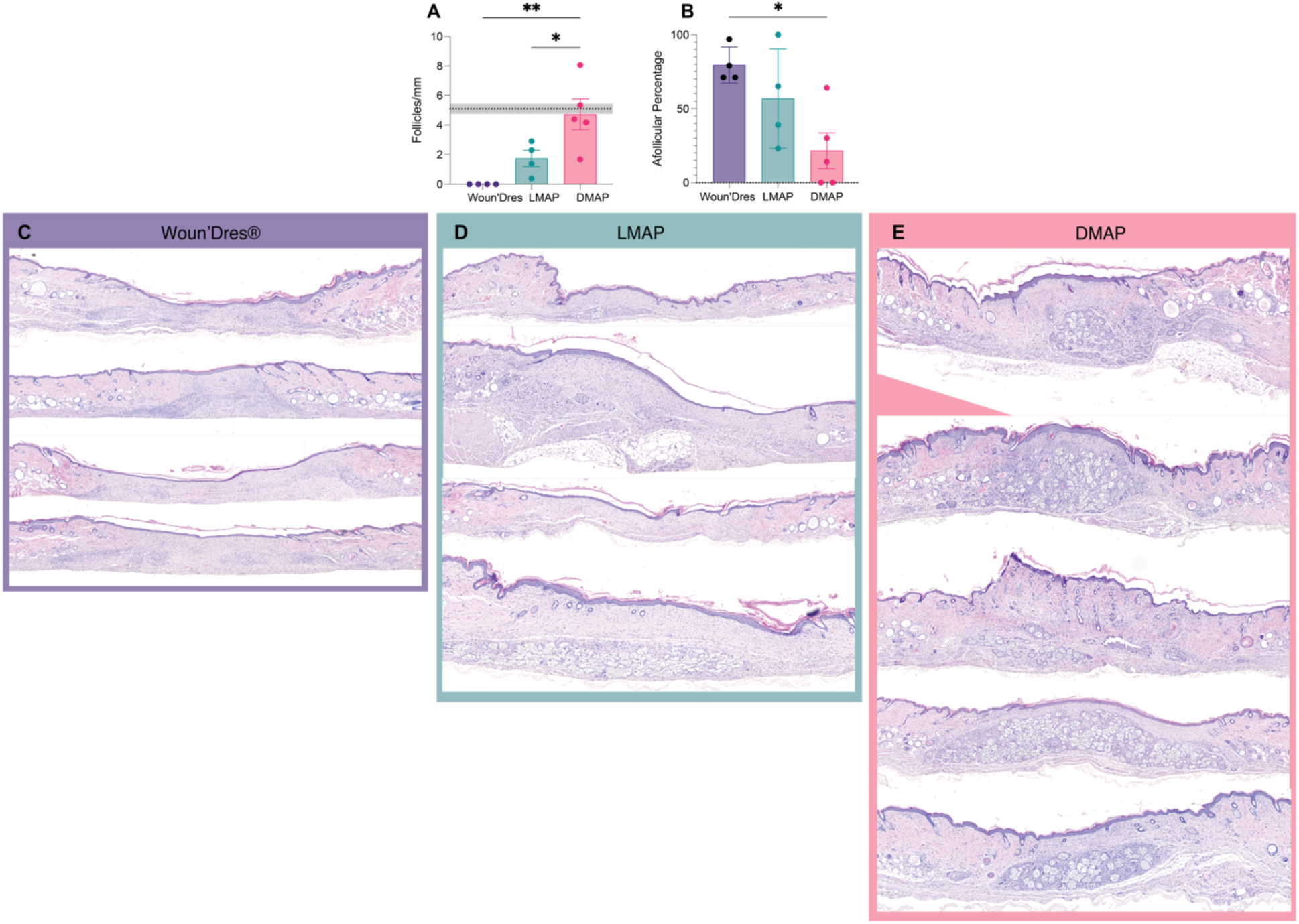
Histological assessment of wound healing outcomes with Woun’Dres®, LMAP, and DMAP treatments at day 21. **A-B)** Quantification of wound healing features at Day 21 post-injury including (A) number of hair follicles per wound normalized to wound length and (B) afollicular percentage (greatest distance between two dermal features divided by overall wound length, indicates true scar). One-way ANOVA with Tukey’s multiple comparisons test was performed. Statistical significance is indicated (*p < 0.05; **p < 0.01). **C–E)** Representative H&E-stained histological sections of healed wounds at Day 21 for (C) Woun’Dres®, **(D)** LMAP, and **(E)** DMAP treatment groups. Images illustrate differences in tissue architecture, follicle regeneration, and cellularity among groups.

**Supplemental Figure 3.**
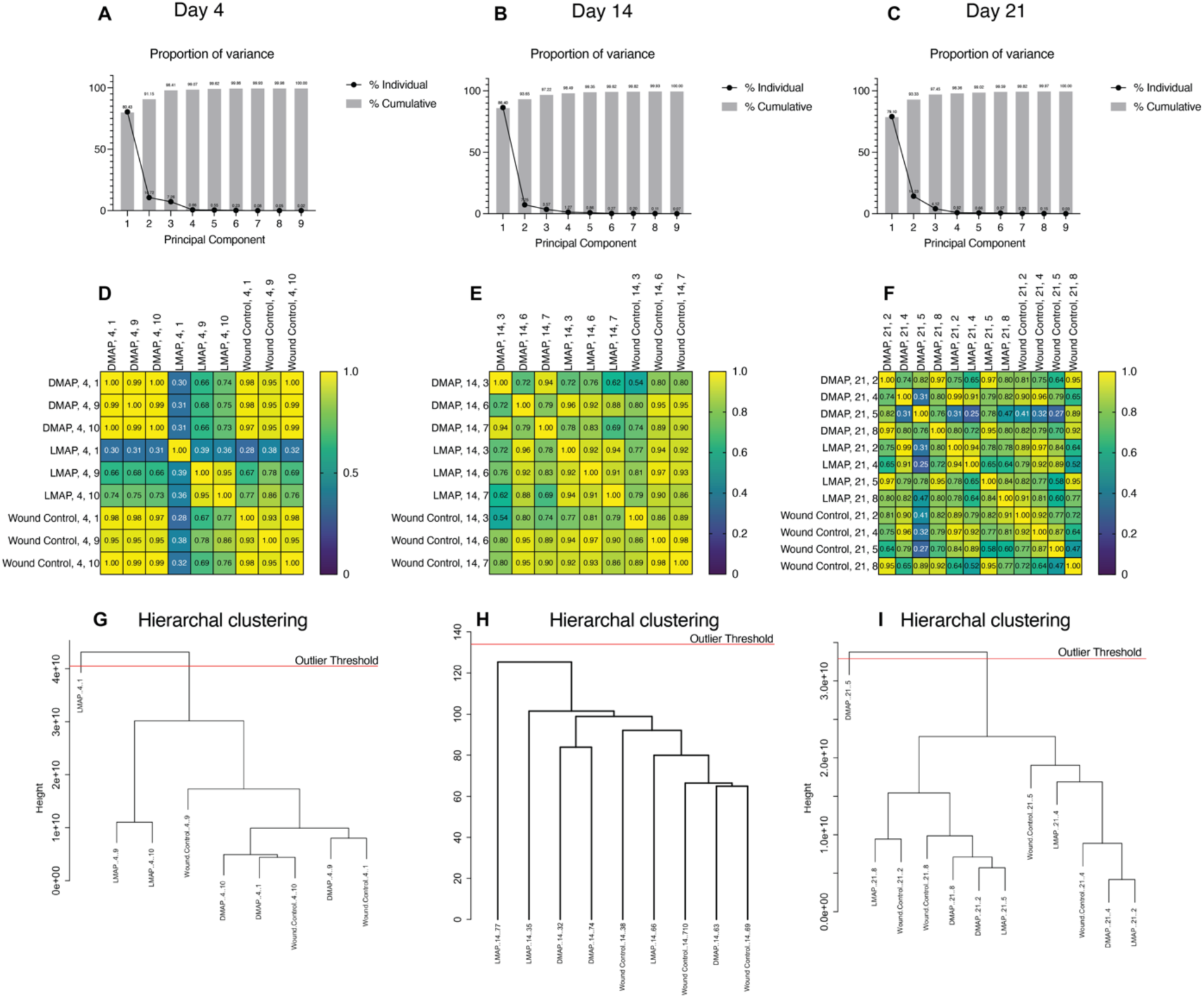
Principal component and cluster analysis of proteomic profiles in wound healing across treatments and timepoints. **A–C)** Principal component analysis (PCA) scree plots showing the proportion of variance explained by each principal component at **(A)** Day 4, **(B)** Day 14, and **(C)** Day 21. The first two principal components capture the majority of variance in the dataset at each timepoint. **D–F**) Pearson correlation plots for all treatment groups (DMAP, LMAP, Woun’Dres®) at **(D)** Day 4, **(E)** Day 14, and **(F)** Day 21. Color scale indicates correlation coefficient (0–1), with higher values representing greater similarity in proteomic profiles. **G–I)** Hierarchical clustering dendrograms for each timepoint: **(G)** Day 4, **(H)** Day 14, and **(I)** Day 21. Clustering reveals the relationships and outlier status among samples from different treatment groups and timepoints, with the outlier threshold indicated by a red line.

**Supplemental Figure 4.**
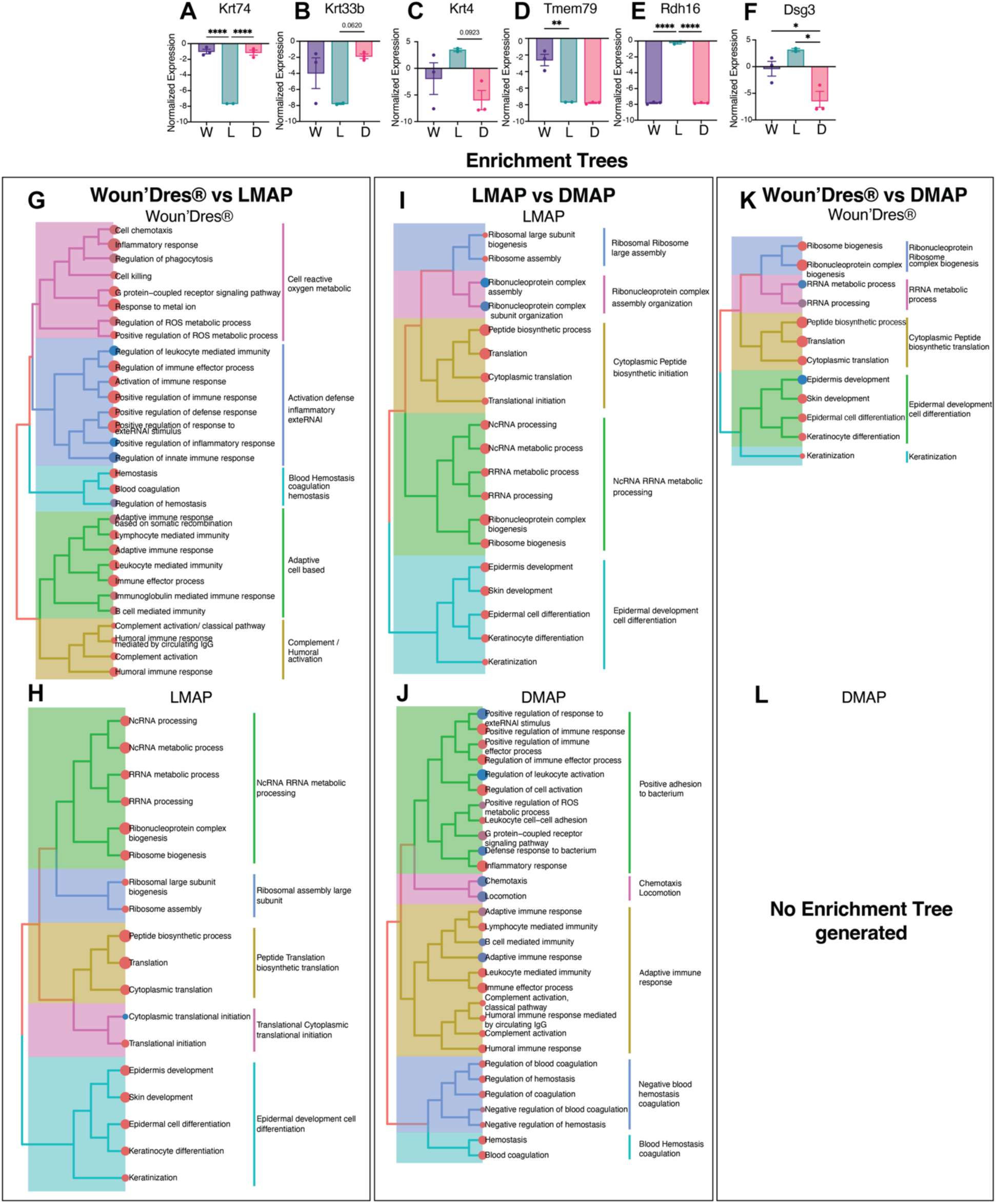
Differential protein expression and pathway enrichment analysis across Woun’Dres®, LMAP, and DMAP treatments at day 4. **A–F)** Bar graphs showing normalized expression levels of selected proteins: **(A)** Krt74, **(B)** Krt33b, **(C)** Krt4, **(D)** Tmem79, **(E)** Rdh16, **(F)** Dsg3 in wound tissue at day 4 across treatment groups: Woun’Dres® (W), LMAP (L), and DMAP (D). One-way ANOVA with Tukey’s multiple comparisons test was performed. Significance is indicated (*p < 0.05, **p < 0.01, ***p < 0.001, ****p < 0.0001). **G–L)** Enrichment trees illustrating hierarchical clustering of significantly enriched biological pathways for each treatment group and pairwise comparison: **G-H)** Woun’Dres® vs LMAP: Pathways upregulated in (**G)** Woun’Dres® and **(H)** LMAP, **I-J)** LMAP vs DMAP: Pathways upregulated in **(I)** LMAP and **(J)** DMAP, **K-L)** Woun’Dres® vs DMAP: Pathways upregulated in **(K)** Woun’Dres® and **(L)** DMAP. Colored branches group related biological processes.

**Supplemental Figure 5.**
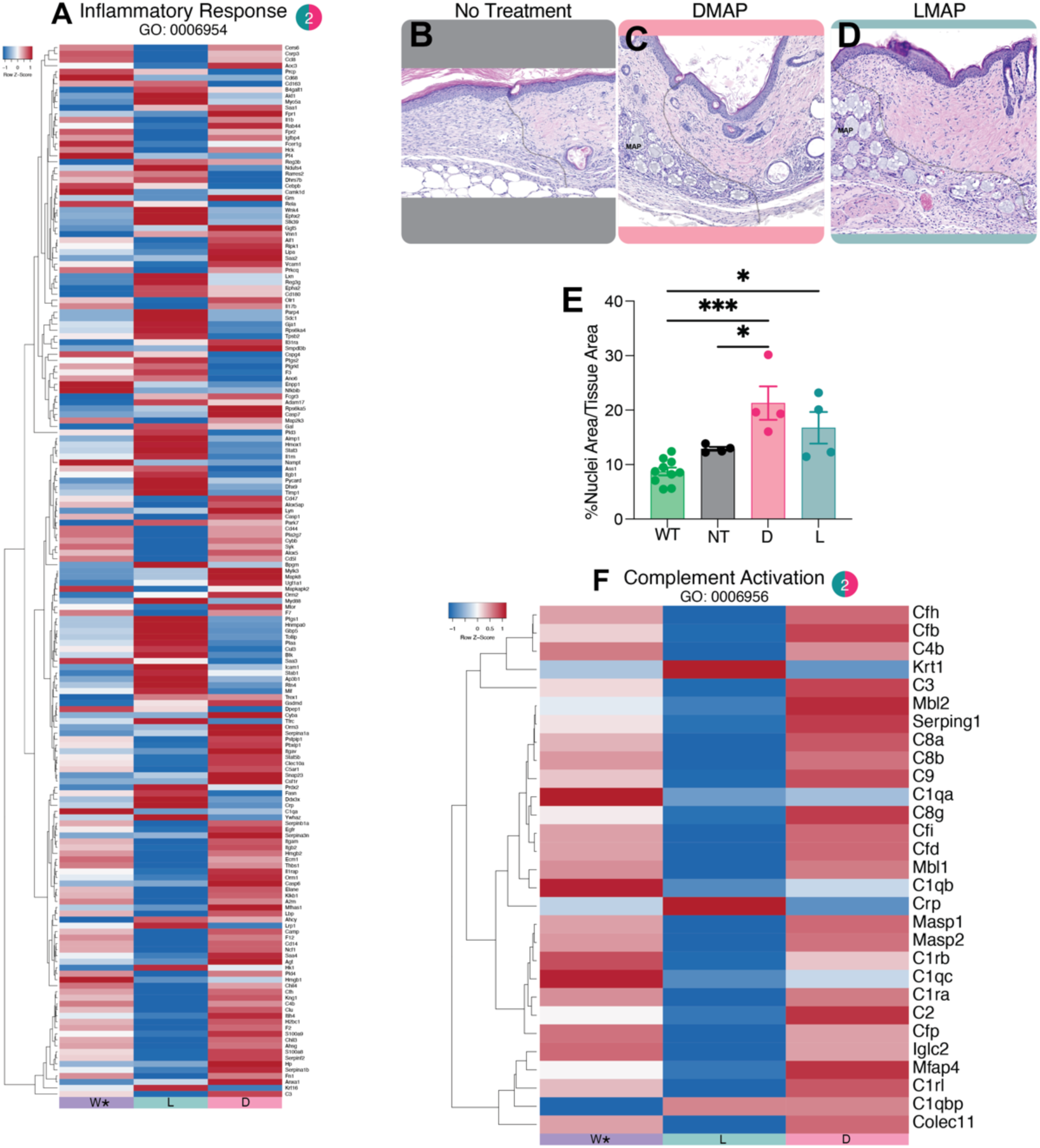
LMAP and DMAP treatments induce distinct inflammatory and complement protein expression signatures and alter skin histology at day 7. **A)** Heatmap showing differential expression of proteins associated with the inflammatory response (Gene Ontology: GO:0006954) across three groups Woun’Dres® (W), LMAP-treated (L), and DMAP-treated (D) skin samples. Each row represents a protein, and each column represents an individual sample. Red indicates upregulation, and blue indicates downregulation relative to the mean expression. **B–D)** Representative H&E-stained histological sections of skin from **(B)** no treatment (NT), **(C)** DMAP-treated, and (**D)** LMAP-treated groups. Scale bar, 100 μm. **(E)** Quantification of nuclear area as a percentage of total tissue area in the outer 20% of the wound (infiltrating edge) in wildtype (WT), NT, DMAP, and LMAP groups. Data are shown as mean ± SEM. Statistical significance was determined by one-way ANOVA with post hoc Tukey tests (*p < 0.05, ***p < 0.001). **F)** Heatmap of proteins involved in complement activation (GO:0006956) across Woun’Dres®, LMAP, and DMAP groups. Red indicates upregulation, and blue indicates downregulation relative to the mean expression.

**Supplemental Figure 6.**
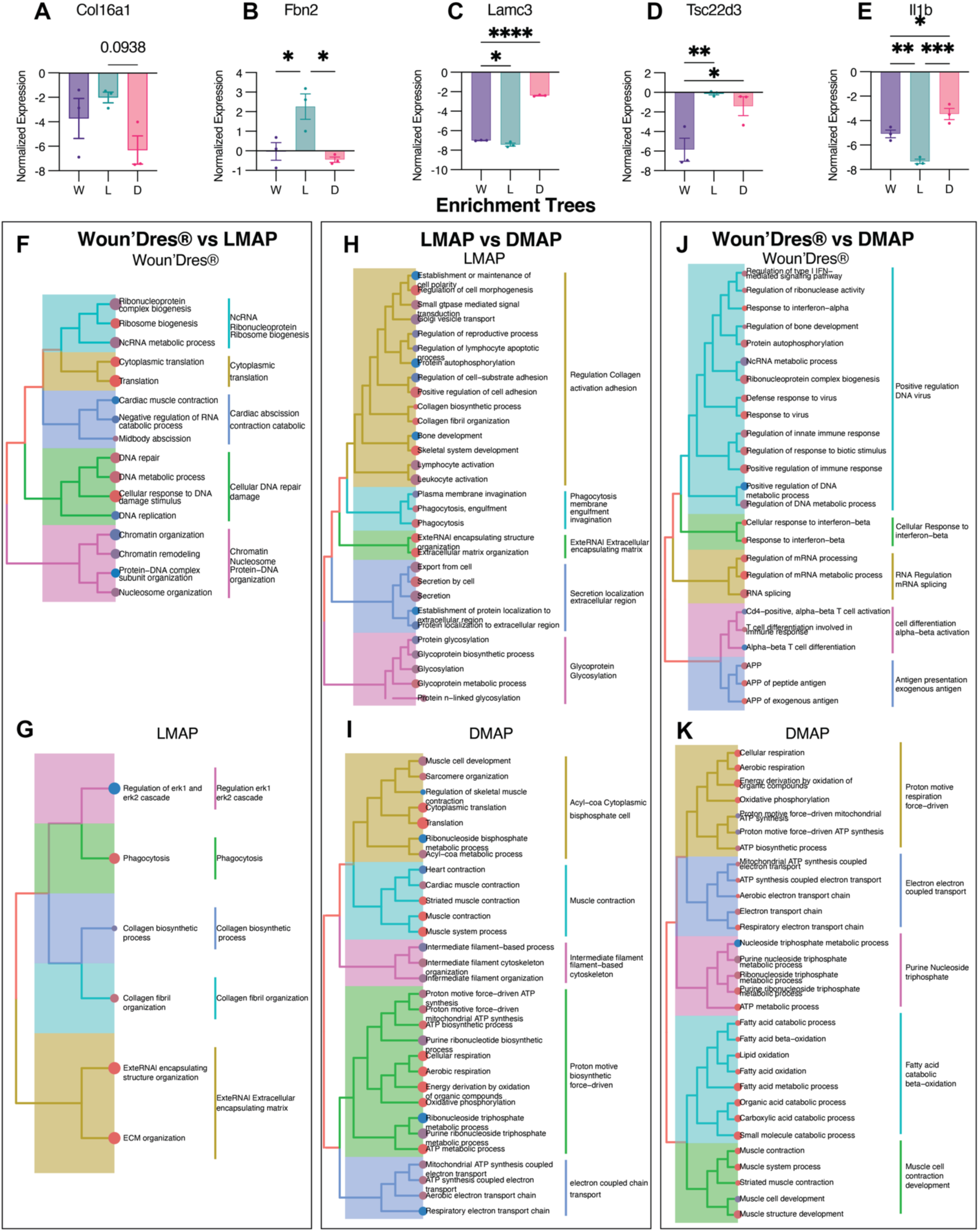
Differential protein expression and pathway enrichment analysis across Woun’Dres®, LMAP, and DMAP treatments at day 14. **A–E)** Bar graphs showing normalized expression levels of selected proteins: **(A)** Col16a1, **(B)** Fbn2, **(C)** Lamc3, **(D)** Tsc22d3, **(E)** Il1b in wound tissue at Day 14 across treatment groups: Woun’Dres® (W), LMAP (L), and DMAP (D). One-way ANOVA with Tukey’s multiple comparisons test was performed. Statistical significance is indicated (*p < 0.05, **p < 0.01, ***p < 0.001, ****p < 0.0001). **F–K)** Enrichment trees illustrating hierarchical clustering of significantly enriched biological pathways for each treatment group and pairwise comparison: **F-G)** Woun’Dres® vs LMAP: Pathways upregulated in **(F)** Woun’Dres® and **(G)** LMAP, **H-I)** LMAP vs DMAP: Pathways upregulated in **(H)** LMAP and **(I)** DMAP, **J-K)** Woun’Dres® vs DMAP: Pathways upregulated in **(J)** Woun’Dres® and **(K)** DMAP. Colored branches group related biological processes.

**Supplemental Figure 7.**
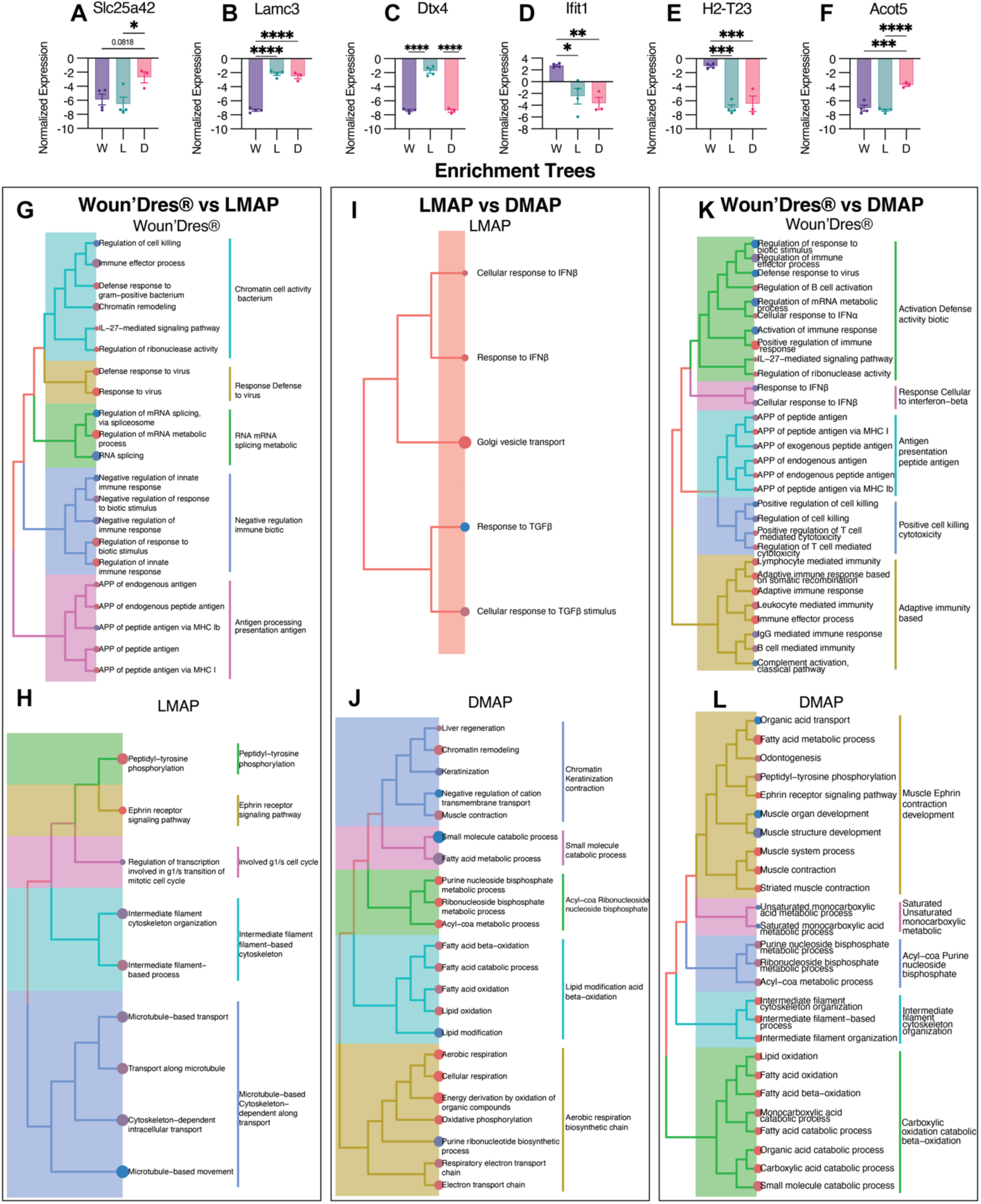
Differential protein expression and pathway enrichment analysis across Woun’Dres®, LMAP, and DMAP treatments at day 21. **A–F)** Bar graphs showing normalized expression levels of selected proteins **(A)** Slc25a2, **(B)** Lamc3, **(C)** Dtx4, **(D)** Ifit1, **(E)** H2-T23, **(F)** Acot5 in wound tissue at Day 21 across treatment groups: Woun’Dres® (W), LMAP (L), and DMAP (D). One-way ANOVA with Tukey’s multiple comparisons test was performed. Statistical significance is indicated (*p < 0.05, **p < 0.01, ***p < 0.001, ****p < 0.0001). **G–L)** Enrichment trees illustrating hierarchical clustering of significantly enriched biological pathways for each treatment group and pairwise comparison: **G-H)** Woun’Dres® vs LMAP: Pathways upregulated in **(G)** Woun’Dres® and **(H)** LMAP, **(I-J)** LMAP vs DMAP: Pathways upregulated in **(I)** LMAP and **(J)**DMAP, **K-L)** Woun’Dres® vs DMAP: Pathways upregulated in **(K)** Woun’Dres® and **(L)** DMAP. Colored branches group related biological processes.

**Supplemental Figure 8.**
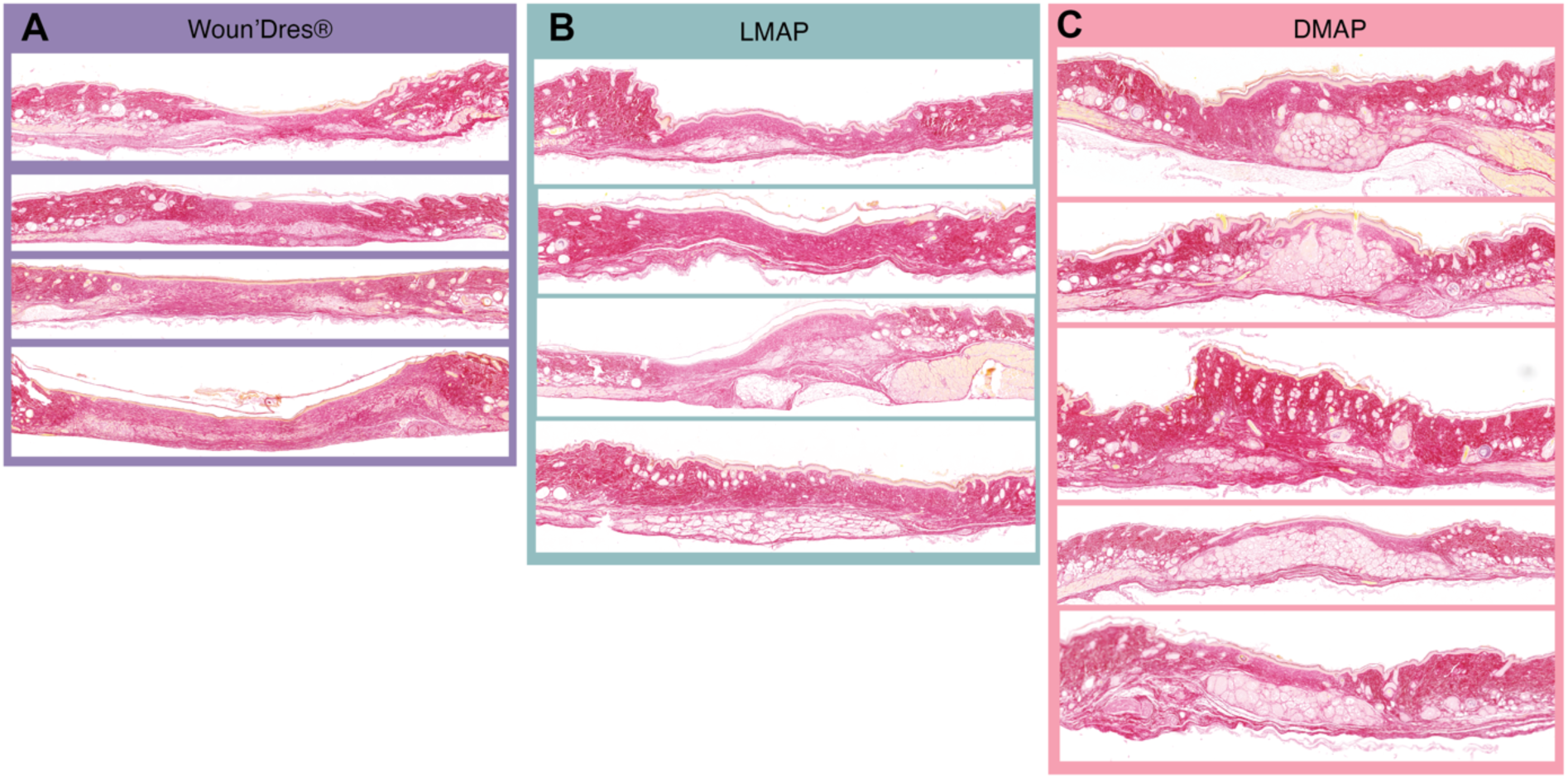
Picrosirius red staining reveals collagen deposition in healed wounds treated with Woun’Dres®, LMAP, or DMAP. **A–C)** Representative Picrosirius Red-stained sections of healed skin wounds from mice treated with **(A)** Woun’Dres®, **(B)** LMAP, and **(C)** DMAP. Multiple sections per group illustrate collagen fiber distribution and organization within the wound bed for each treatment. Collagen appears red, highlighting differences in extracellular matrix remodeling among groups.

## Notes

### Competing Interest Statement

T.S. is a founder of Tempo Therapeutics which aims to commercialize MAP technology.

